# Sequence-encoded H2A.Z nucleosome dynamics control DNA unwrapping and SUV420H1 recognition

**DOI:** 10.64898/2026.06.10.731474

**Authors:** Tong Zhang, Li Huang, Xuanfeng Li, Bingyan Liu, Juan Li, Chaowei Shi, Shusheng Fu, Zheng Zhou, ShengQi Xiang

**Author notes:** These authors contributed equally to this work (Tong Zhang, Li Huang).

## Abstract

H2A.Z and canonical H2A adopt nearly identical nucleosomal folds, yet their distinct chromatin functions are not captured by static structural analysis. Using fast magic-angle spinning ^1^H-detected solid-state NMR, we show that H2A.Z possesses enhanced backbone flexibility in the L1 loop and the α2–L2 region (M2) relative to H2A. Chimeric segment-swapping demonstrates that these dynamic signatures are locally sequence-encoded and functionally transplantable. The inherent mobility of the M2 region promotes nucleosomal DNA-end unwrapping and persists when DNA ends are stabilized by linker histone H1 or opened by SUV420H1, indicating that this mobility is intrinsic rather than a passive consequence of DNA detachment. Chemical shift perturbation mapping and catalytic assays further show that SUV420H1 reads this H2A.Z-specific conformational landscape: the M2 region, together with the H2A.Z DS motif, supports variant-selective methyltransferase activity. These findings establish an axis of sequence-dynamics-accessibility-recognition along which local backbone fluctuations serve as physical determinants of epigenetic enzyme specificity.

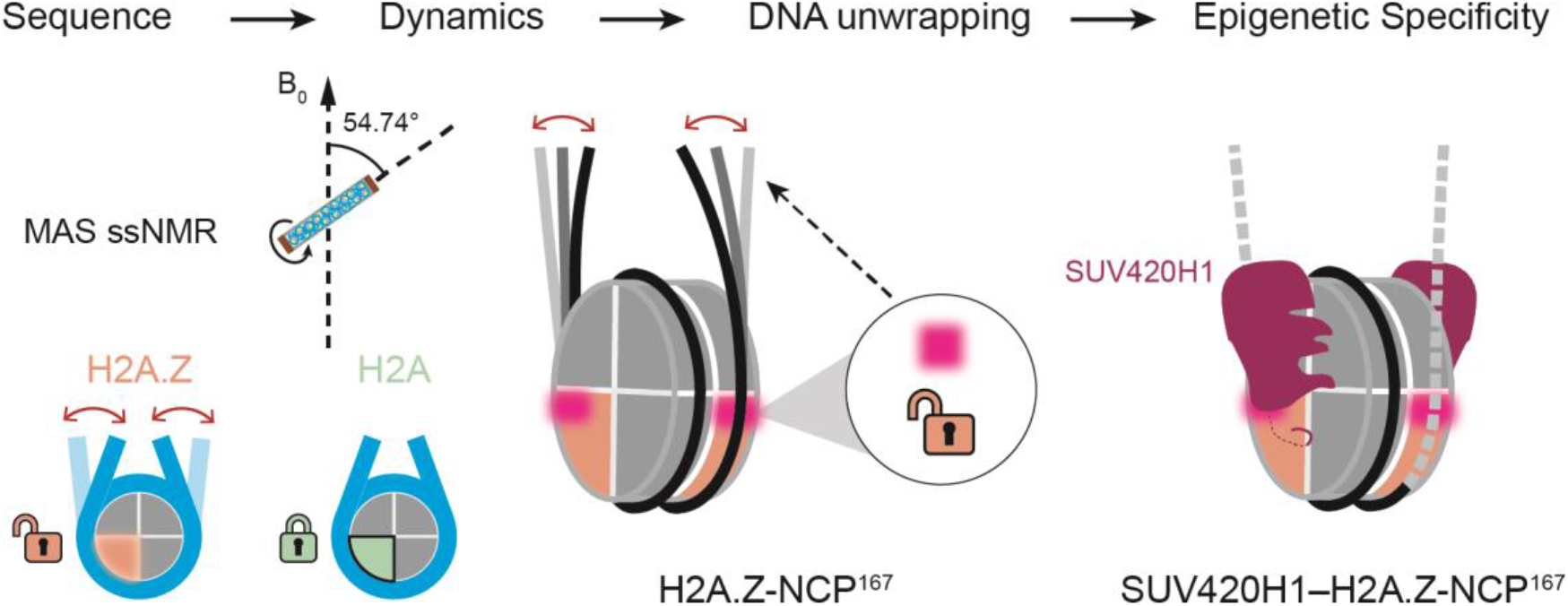

## Introduction

In eukaryotes, genomic DNA is compacted into chromatin, whose fundamental repeating unit, the nucleosome core particle (NCP), consists of ~147 bp of DNA wrapped around a histone octamer (H2A, H2B, H3, H4)^1–4^. Far from static, nucleosomes undergo dynamic behaviors including DNA unwrapping, sliding, and conformational fluctuations within the histone core, processes critical for regulating DNA accessibility^3,5–9^.

The functional diversity of chromatin is substantially expanded by histone variants, non-allelic isoforms that replace canonical counterparts in a targeted, replication-independent manner^10–14^. Among these, H2A.Z is a highly conserved H2A variant enriched at promoters, enhancers, and replication origins, where it participates in transcription, DNA repair, and replication^15–28^.

Despite their well-established functional divergence, H2A.Z and canonical H2A adopt nearly identical nucleosomal structures^29^. High-resolution structures reveal only subtle differences in the L1 loop, acidic patch, and docking domain, yet whether these confer stabilizing or destabilizing effects remains controversial^16,29–38^. This ambiguity reflects an inherent limitation of static methods: crystallography and cryo-EM provide exquisite architectural snapshots but cannot capture the intrinsic backbone flexibility and conformational fluctuations that distinguish highly similar macromolecular complexes. Indeed, H2A.Z nucleosomes exhibit increased micrococcal nuclease (MNase) sensitivity and enhanced DNA end accessibility^35–38^, suggesting that dynamic properties, which are potentially encoded by local sequence differences, may underlie their functional specificity^35,37–41^. However, a residue-level understanding of how H2A.Z modulates nucleosome dynamics, and how such dynamics are “read” by downstream effectors such as SUV420H1, has remained elusive. Magic-angle spinning (MAS) solid-state NMR (ssNMR) spectroscopy has emerged as a powerful tool for probing the dynamics of macromolecular complexes at atomic resolution^42–48^. Both ^1^H-detected and ^13^C-detected ssNMR approaches have enabled high-quality resonance assignments of histones within the nucleosome, facilitating subsequent studies of interactions and dynamics^43,45,48–53^.

In contrast to the relatively well-characterized mechanisms by which H2A.Z–H2B dimers are selectively recognized by histone chaperones and remodelers, how H2A.Z is preferentially recognized within the intact nucleosome context has remained less understood^54^. H2A.Z facilitates early replication origin activation through preferential recognition by the methyltransferase SUV420H1, which deposits H4K20me2 to enable ORC1 binding^22^. Recent cryo-EM studies provided, to our knowledge, the first structural framework for variant-selective recognition of H2A.Z-containing nucleosomes by SUV420H1^55^, revealing that SUV420H1 engages the nucleosome through contacts with the H4 tail, DNA, and the acidic patch, and that its KR loop interacts with H2A.Z-specific D97/S98 residues (the DS motif), to promote H2A.Z-nucleosome preference^55–57^. However, reciprocal exchange of the DS/NK motif between H2A.Z and H2A does not fully interchange SUV420H1 activity toward the two nucleosome substrates, indicating that elements beyond the DS motif contribute to variant preference. This incomplete conversion may reflect an incompletely mapped SUV420H1– nucleosome interaction landscape, or alternatively, an additional contribution from intrinsic H2A.Z nucleosome dynamics. Thus, although static structures have established an important framework for SUV420H1 recognition of H2A.Z-containing nucleosomes, how intrinsic H2A.Z dynamics contribute to this preferential recognition remains unresolved.

In this study, we employed high-resolution ssNMR spectroscopy to dissect the intrinsic conformational dynamics of H2A.Z and H2A within the nucleosome environment. Through comprehensive backbone resonance assignments and site-specific dynamic analysis of both variants in this study, we uncovered two H2A.Z-specific dynamic regions: the L1 loop and a previously uncharacterized α2–L2 region, hereafter termed M2. Using a series of segment-swapped chimeric mutants, we demonstrate that this flexibility is locally encoded by amino acid sequence and thus intrinsic to H2A.Z. We further show that the M2 dynamic region promotes DNA-end unwrapping, persists under H1- and SUV420H1-bound chromatin states, and contributes to SUV420H1 variant-selective recognition and catalysis. Finally, by integrating chemical shift perturbation (CSP) mapping with Y349 mutagenesis, we distinguish H2A.Z-selective dynamic recognition from a conserved catalytic engagement interface. Together, these results provide a mechanistic framework in which H2A.Z-specific local dynamics are converted into DNA accessibility and epigenetic enzyme specificity.

## Results

### H2A.Z nucleosomes contain sequence-encoded dynamic regions

The ssNMR samples of H2A.Z and H2A nucleosomes were prepared as previously described^58^, yielding transparent gels with concentrations consistent with the typical chromatin concentration in cells^59^, ranging from approximately 100 to 200 mg/mL(Figure 1A; Figure S1A–S1C; Figure S4A). Well-resolved two- and three-dimensional (2D/3D) correlation spectra were acquired using ^1^H-detected ssNMR experiments at 60 kHz MAS, achieving resolution comparable to previous reports^49^. For H2A.Z, assignments of the rigid core (residues 14–118) were obtained by simultaneously matching Cα and CO chemical shifts across a set of proton-detected triple-resonance experiments, including CANH, CA(CO)NH, CONH, and CO(CA)NH spectra (Figure 1A–1C), with amino acid types identified by the Cβ chemical shifts from a 3D CBCANH spectrum (Figure 1D) employing the PC4 sequence^60^. For H2A, backbone assignments were transferred from Drosophila nucleosomes^49^ with the help of CANH, CA(CO)NH, and CONH (Figure S4A, B). In total, assignments were completed for 87 of 102 residues (85%) in H2A.Z (14–118) and 92 of 94 (98%) in H2A (16–113) (Figure 1A; Figure S2B; Figure S4A). As shown in Figure S2A, S2C, and S4C, secondary structure analysis showed that their structures match the corresponding cryo-EM and X-ray structures (PDB: 8THU^56^ for H2A.Z and PDB: 5GT3^61^ for H2A).

**Figure 1.**
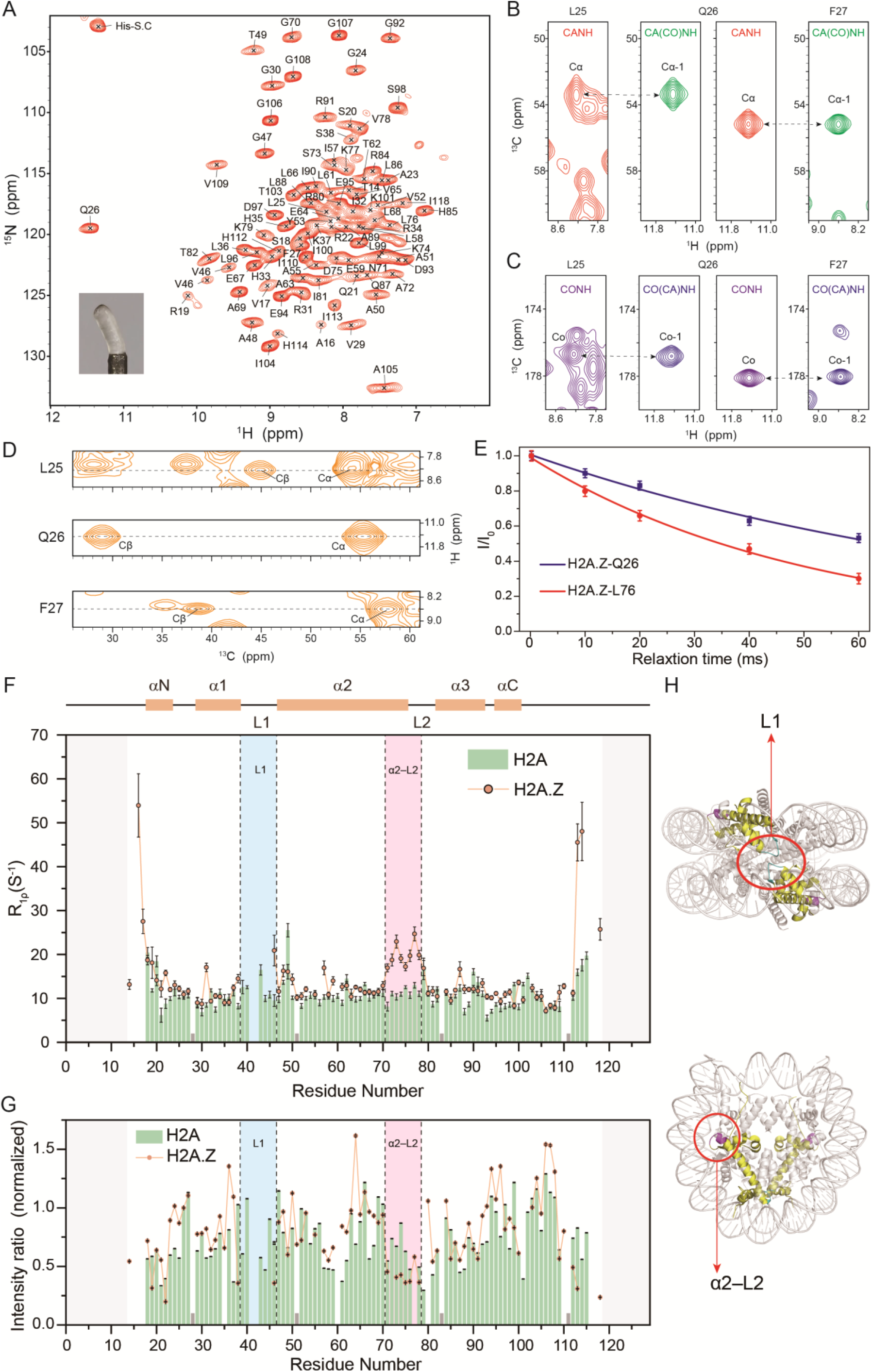
Backbone resonance assignment and distinct dynamic features of H2A.Z in nucleosomes. (A) 2D NH spectrum of H2A.Z in sedimented nucleosomes (pH 6.5). A total of 87 assigned residues are indicated. Inset: photograph of the gel-like sedimented nucleosome sample used for solid-state NMR experiments. (B–D) Representative strips for residues L25→Q26→F27 illustrating the sequential backbone assignment of H2A.Z. (B) Cα correlations from 3D CANH (red) and CA(CO)NH (green) spectra. (C) CO correlations from 3D CONH (purple) and CO(CA)NH (blue) spectra. (D) Cβ correlations from the 3D CBCANH spectrum (yellow), aiding amino acid type identification and validating assignment accuracy. (E) Normalized *R*_*1ρ*_ signal intensity decay curves for representative residues Q26 (rigid core) and L76 (α2–L2 region) of H2A.Z, revealing markedly different relaxation profiles. Error bars were calculated based on the signal-to-noise ratio of individual cross-peaks. (F) Site-specific ^15^N *R*_*1ρ*_ profiles comparing H2A.Z and H2A. The α2–L2 region (from N71 to V78) exhibits markedly elevated *R*_*1ρ*_ values in H2A.Z, indicating enhanced backbone flexibility. The secondary structure of H2A.Z in the nucleosome (PDB:8THU^56^) is shown at the top. Gray bars denote proline residues. (G) Normalized signal intensity comparison from 3D CANH spectra (H2A.Z vs. H2A). Residues in the H2A.Z α2–L2 region (from N71 to V78) display markedly lower intensities compared to other H2A.Z domains and the corresponding region in H2A, corroborating its enhanced flexibility. (H) Structural mapping of the dynamically distinct regions (L1 and α2–L2) in H2A.Z and H2A within the nucleosome.

Despite structural similarity, H2A.Z and H2A exhibited markedly different dynamic profiles. We measured ^15^N *R*_*1ρ*_ relaxation rates for both H2A.Z and H2A within nucleosomes under 60 kHz MAS with a 9 kHz spin-lock field (Figure 1E, F). In the H2A.Z nucleosome, residues in the L1 loop (from R39 to R45) were undetectable in cross-polarization (CP) spectra, and V46 showed a large *R*_*1ρ*_ value, suggesting substantial motion (Figure 1F). Notably, residues in the region encompassing the C-terminal part of the α2 helix and the adjacent part of L2 loop (α2–L2, from N71 to V78) displayed significantly elevated *R*_*1ρ*_ values compared to both the other rigid regions of H2A.Z and the corresponding region in H2A (Figure 1E, F). This indicates a H2A.Z-specific backbone flexibility in the α2–L2 region (residues N71–V78). This enhanced flexibility was corroborated by reduced signal intensities in 3D CANH spectra for residues N71 to V78 relative to other sites in H2A.Z, a contrast absent in the corresponding region of H2A (Figure 1G).

To test whether these distinct dynamic properties are determined by local sequence differences, we designed a series of chimeric mutants by swapping regions that encompass the dynamic segments plus flanking sequence-variable residues between H2A.Z and H2A (Figure 2A). Specifically, we generated three chimeric mutants: H2A.Z-M1, in which the M1 region of H2A.Z (residues K37–V46) was replaced with that of H2A (residues R35–V43); H2A.Z-M2, in which the M2 region of H2A.Z (residues N71–T82) was replaced with that of H2A (residues N68–I79); and H2A-M2, in which the corresponding region of H2A (residues N68–I79) was replaced with that of H2A.Z (residues N71–T82). All mutants successfully assembled into histone octamers and nucleosomes (Figure S1D, E).

**Figure 2.**
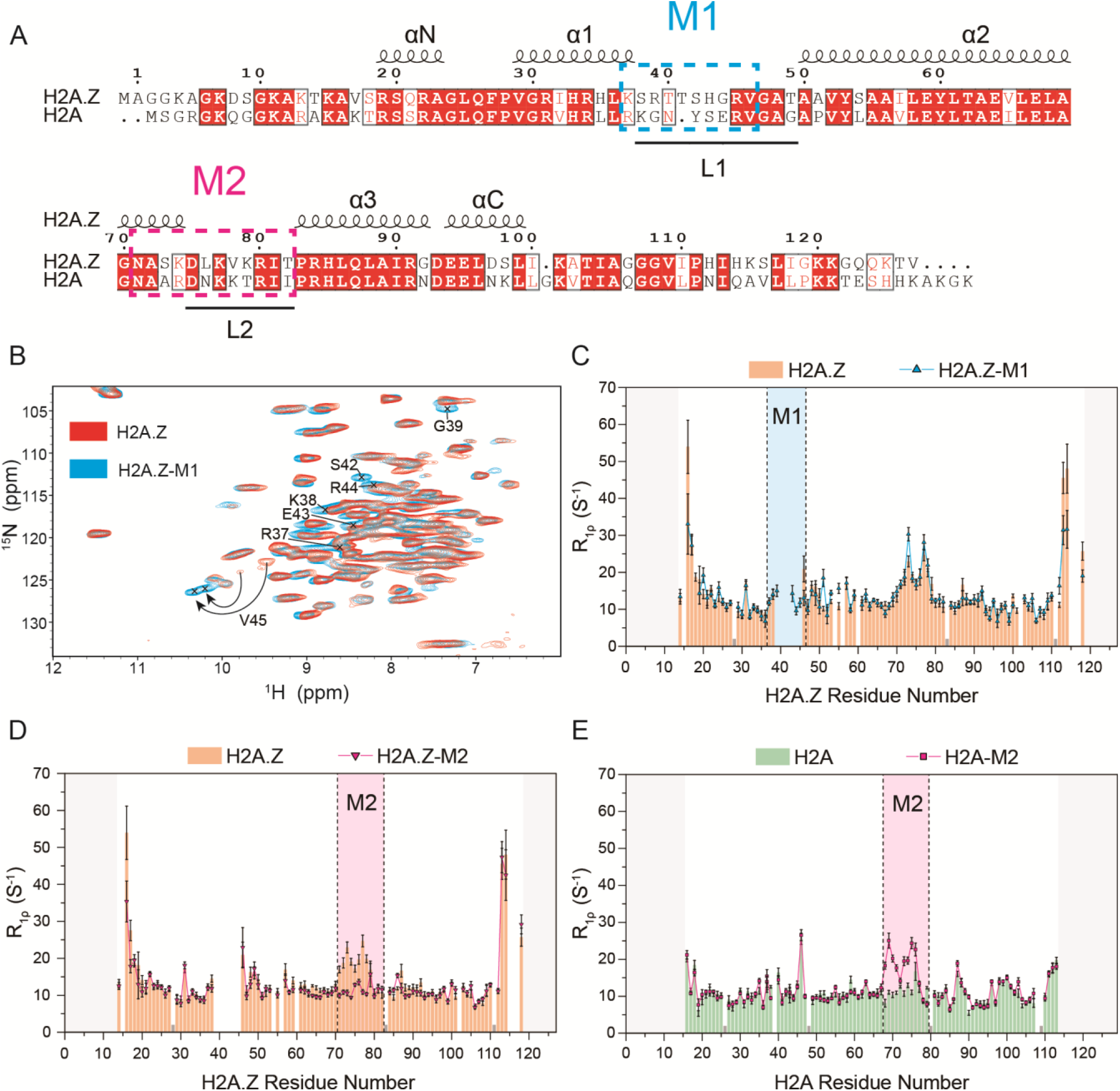
Sequence-swapping experiments demonstrate that regional dynamic differences between H2A.Z and H2A are encoded by local amino acid sequences. (A) Sequence alignment of H2A.Z and H2A. Identical residues are shown as white characters on a red background; similar residues are shown in red and boxed. Dashed boxes indicate the positions of the mutated regions M1 (cyan) and M2 (pink), which encompass the dynamic difference regions L1 and α2–L2, respectively. (B) Overlay of 2D NH correlation spectra (H2A.Z vs. H2A.Z-M1). Replacement of the H2A.Z M1 region with the corresponding H2A sequence rendered the L1 loop residues (G39, S42, E43, R44) detectable in cross-polarization spectra, whereas these resonances are absent in wild-type H2A.Z due to enhanced mobility. (C) Site-specific ^15^N *R*_*1ρ*_ profiles comparing H2A.Z and H2A.Z-M1. In H2A.Z-M1, *R*_*1ρ*_ values in the M1 region return to baseline levels, confirming attenuation of M1 dynamics. (D) Site-specific ^15^N *R*_*1ρ*_ profiles comparing H2A.Z and H2A.Z-M2. In H2A.Z-M2, the elevated M2 region *R*_*1ρ*_ values observed in wild-type H2A.Z are abolished, returning to baseline, indicating the H2A-derived sequence attenuates flexibility. (E) Site-specific ^15^N *R*_*1ρ*_ profiles comparing H2A and H2A-M2. In H2A-M2, the M2 region exhibits a pronounced increase in *R*_*1ρ*_ values, indicating that the H2A.Z-derived M2 sequence confers enhanced flexibility into H2A. Gray bars in (C), (D), and (E) denote proline residues.

^15^N *R*_*1ρ*_ measurements on these mutant nucleosomes under the same conditions as wild type revealed that dynamic features are transferable with the sequence (Figure 2B–2E; Figure S3; Figure S5). In H2A.Z-M1, the previously invisible L1 residues in wild-type (WT) H2A.Z became detectable in CP spectra, and their *R*_*1ρ*_ values dropped to baseline levels (Figure 2B, C), indicating successful reversal of M1 region dynamics. Data from the M2 region further supported this sequence-dependent origin of differential dynamics. In H2A.Z-M2, the *R*_*1ρ*_ values in the swapped region decreased to the rigid core baseline, effectively eliminating the elevated flexibility seen in WT H2A.Z (Figure 2D). Conversely, in H2A-M2, this region exhibited markedly increased *R*_*1ρ*_, demonstrating that the characteristic flexibility of the H2A.Z M2 region was successfully introduced into H2A (Figure 2E). These results demonstrate that the distinct backbone dynamics of H2A.Z and H2A are determined by their local amino acid sequences.

### The H2A.Z M2 dynamic region promotes nucleosomal DNA-end unwrapping

Given the spatial proximity of the M2 region to the DNA entry-exit site on the nucleosome (Figure 1H), we hypothesized that its enhanced mobility might facilitate local DNA unwrapping. To examine whether enhanced mobility correlated with increased DNA accessibility, we first compared DNA accessibility using MNase digestion assays (Figure 3A). DNA within H2A.Z nucleosomes was digested more rapidly than that within H2A nucleosomes, as evidenced by a lower fraction of protected 147 bp DNA after MNase digestion (Figure 3B, C), consistent with previous reports linking H2A.Z to increased DNA accessibility^36,38^. Critically, both M2 swap mutants exhibited intermediate nucleosome stability in MNase digestion assays (H2A.Z < chimeras < H2A): H2A.Z-M2, carrying the rigid H2A M2 sequence, showed greater resistance to MNase digestion than WT H2A.Z, whereas H2A-M2, carrying the flexible H2A.Z M2 sequence, showed increased susceptibility to MNase cleavage (Figure 3B, C). These opposing shifts indicate that M2 dynamics directly govern nucleosomal DNA-end unwrapping, increasing local DNA accessibility at the end-exit site.

**Figure 3.**
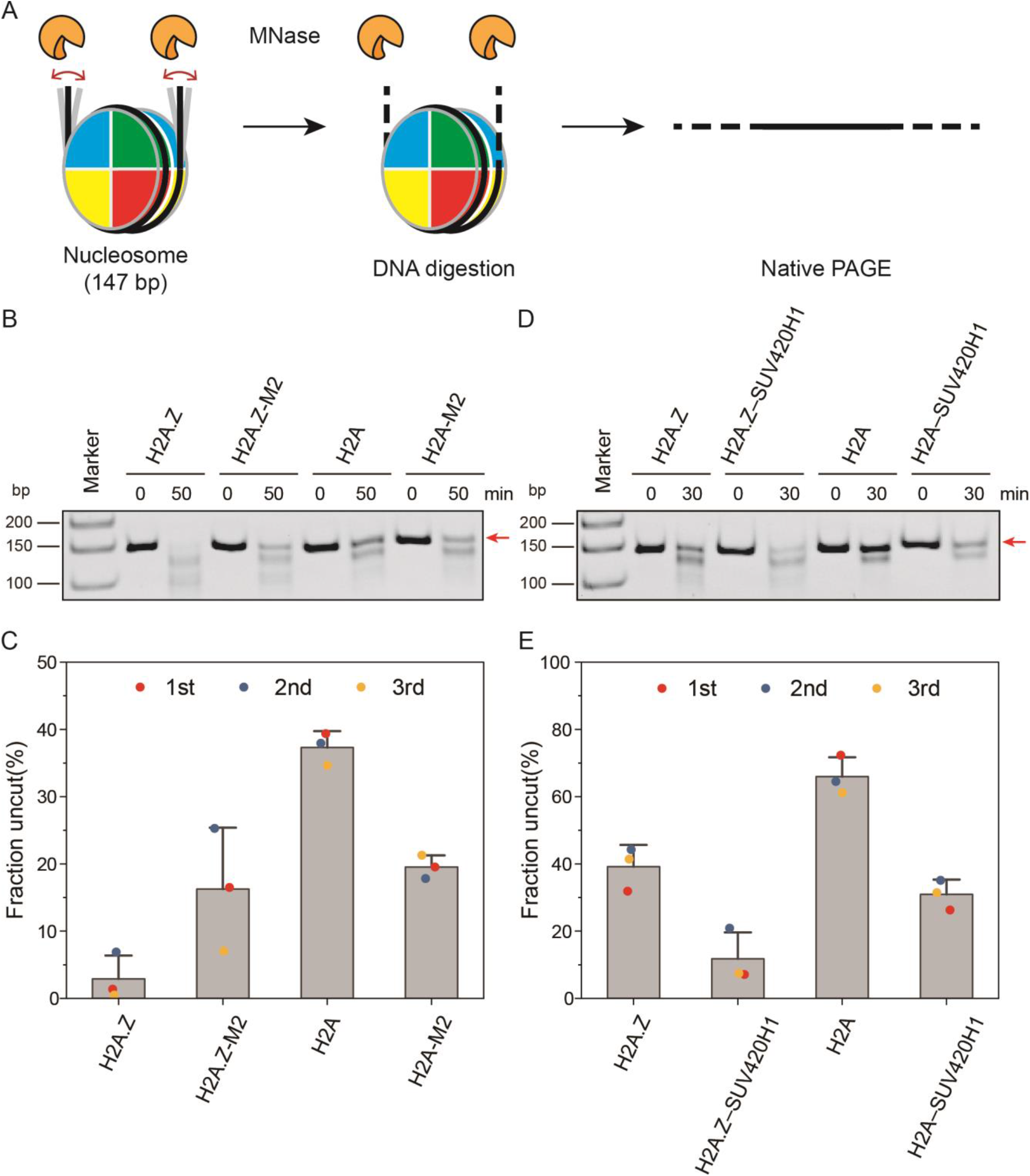
M2 region dynamics and SUV420H1 binding regulate nucleosomal DNA end stability. (A) Schematic of the micrococcal nuclease (MNase) digestion assay. MNase preferentially cleaves nucleosomal DNA ends. Digestion products were analyzed by native PAGE to assess DNA stability. (B) Representative native PAGE gel showing MNase digestion time courses (0 and 50 min) for H2A.Z, H2A.Z-M2, H2A, and H2A-M2 nucleosomes. After 50 min at 37°C, H2A nucleosomes retain the most uncleaved 147 bp DNA, H2A.Z nucleosomes retain the least, and the mutants H2A.Z-M2 and H2A-M2 show intermediate retention. (C) Quantification of MNase digestion results. Values represent the ratio of uncut 147 bp DNA band intensity after digestion to initial intensity (mean + s.d., n = 3). DNA stability follows the order H2A > H2A-M2 ≈ H2A.Z-M2 > H2A.Z, consistent with the M2 backbone flexibility profile. (D) Representative native PAGE gel showing MNase digestion (0 and 30 min) for free and SUV420H1-bound nucleosomes (H2A.Z, H2A.Z–SUV420H1, H2A, H2A–SUV420H1). After 30 min at 37°C, SUV420H1-bound nucleosomes show considerably less residual 147 bp DNA compared to their free counterparts, indicating that SUV420H1 binding reduces DNA end stability. (E) Quantification of MNase digestion for free and SUV420H1-bound nucleosomes. Values represent the ratio of uncut 147 bp DNA band intensity after digestion to initial intensity (mean + s.d., n = 3). SUV420H1-bound nucleosomes exhibit substantially reduced DNA retention, indicating that SUV420H1 binding promotes DNA end unwrapping.

To determine whether SUV420H1 enzyme binding also alters DNA-end stability, we also compared free and SUV420H1-bound nucleosomes. Nucleosomes bound to SUV420H1 exhibited enhanced susceptibility to enzymatic cleavage compared to free nucleosomes (Figure 3D, E), consistent with previous structural observations that SUV420H1 promotes nucleosomal DNA-end detachment^55,56^. Thus, H2A.Z M2 dynamics and SUV420H1 binding converge on DNA-end opening, suggesting that H2A.Z pre-configures nucleosomes toward a conformation favorable for enzyme engagement.

### Chromatin-factor binding rigidifies H2A nucleosomes but does not erase intrinsic H2A.Z M2 dynamics

To further dissect the causal relationship between M2 region flexibility and DNA unwrapping, we examined whether the flexibility is a cause or a consequence of DNA unwrapping. We therefore assembled NCP^167^ complexes with either linker histone H1.4 or SUV420H1 (Figure 4A, B; Figure S1B, C). H1 is known to stabilize the DNA entry-exit region of nucleosomes by engaging linker DNA and restricting DNA-end breathing^62–67^, whereas SUV420H1 has been reported to promote nucleosomal DNA-end detachment upon binding^55,56^, a result which is validated by MNase digestion assays (Figure 3D, E).

**Figure 4.**
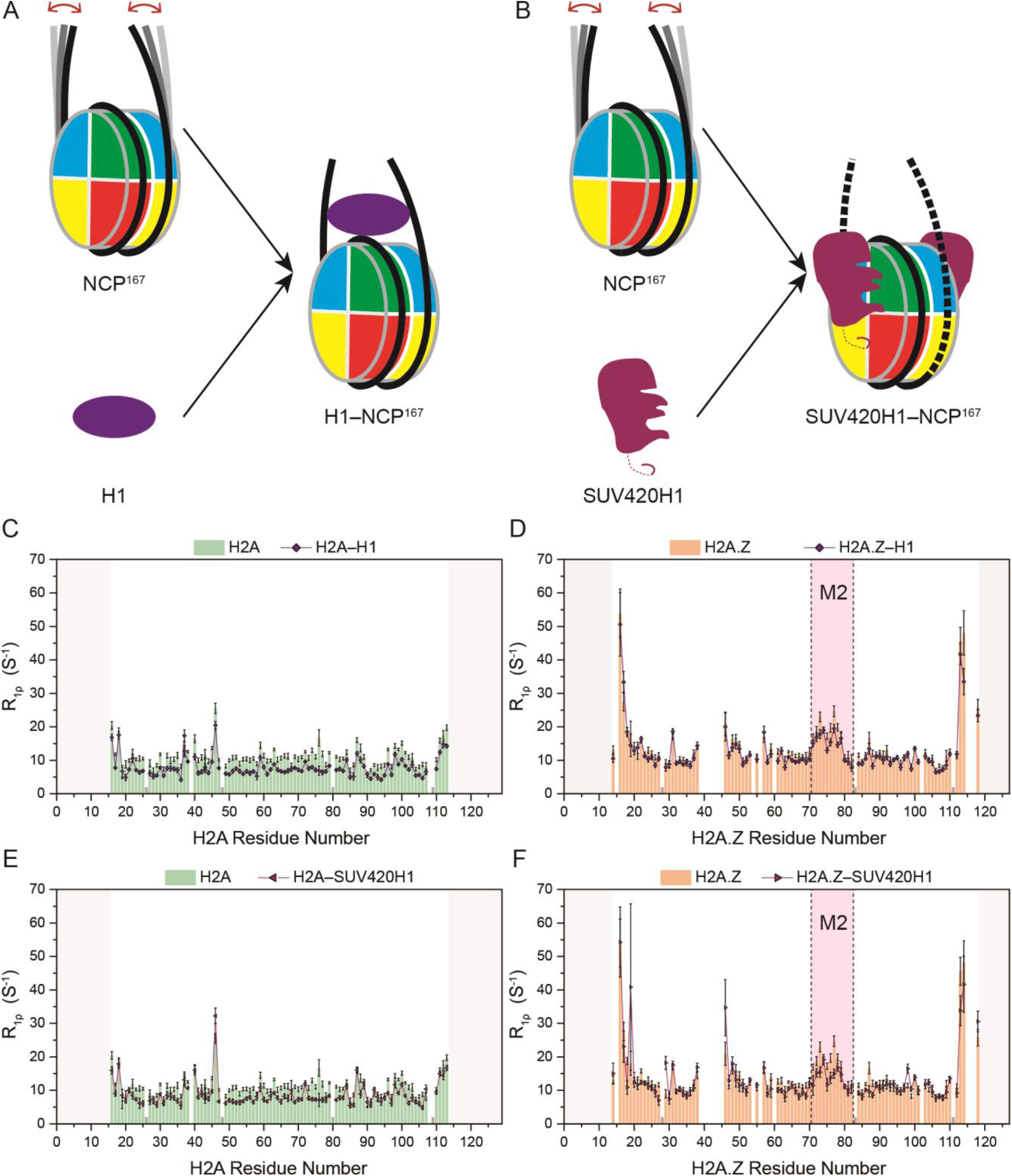
The intrinsic backbone dynamics of the H2A.Z M2 region are independent of nucleosomal DNA unwrapping state. (A) Schematic of H1 binding to the NCP^167^. H1 stabilizes the nucleosome by binding and locking the DNA entry-exit site. (B) Schematic of SUV420H1 binding to the NCP^167^. SUV420H1 promotes nucleosomal DNA end unwrapping. (C) Site-specific ^15^N *R*_*1ρ*_ profiles comparing H2A in the absence and presence of H1. H1 binding causes a global decrease in *R*_*1ρ*_ values, indicating widespread suppression of backbone dynamics. (D) Site-specific ^15^N *R*_*1ρ*_ profiles comparing H2A.Z in the absence and presence of H1. H1 binding does not substantially reduce *R*_*1ρ*_ values in the H2A.Z M2 region; the enhanced flexibility persists. (E) Site-specific ^15^N *R*_*1ρ*_ profiles comparing H2A in the absence and presence of SUV420H1. SUV420H1 binding induces a global decrease in *R*_*1ρ*_ values. (F) Site-specific ^15^N *R*_*1ρ*_ profiles comparing H2A.Z in the absence and presence of SUV420H1. SUV420H1 binding does not substantially alter the *R*_*1ρ*_ values in the M2 region of H2A.Z. Collectively, (C–F) demonstrate that the enhanced flexibility of the H2A.Z M2 region persists regardless of whether DNA ends are stabilized by H1 or unwrapped by SUV420H1, establishing this dynamic feature as an intrinsic, sequence-encoded property.

The ssNMR relaxation data revealed a variant-specific response to chromatin-factor binding. In H2A-containing nucleosomes, both H1.4 and SUV420H1 induced a broad reduction in *R*_*1ρ*_ values across the histone fold, reflecting global rigidification of the canonical nucleosome upon factor binding. (Figure 4C, E; Figure S6A, B; Figure S7A). In contrast, H2A.Z-containing nucleosomes showed only limited global changes upon binding either factor, and the elevated *R*_*1ρ*_ values in the M2 region persisted in both the H1.4- and SUV420H1-bound states (Figure 4D, F; Figure S6C, D; Figure S7B). Thus, the H2A.Z M2 dynamic signature is not erased by a DNA-end-stabilizing factor or by an enzyme associated with DNA-end opening. Together with the M2-swap and MNase results, these data support the conclusion that M2 flexibility is not merely a passive consequence of DNA-end detachment. Rather, it represents an intrinsic, sequence-encoded dynamic property of H2A.Z that remains robust under distinct chromatin-factor perturbations and correlates with increased DNA-end accessibility.

### SUV420H1 reads the H2A.Z M2 dynamic element to promote variant-selective catalysis

Building upon previous insights into SUV420H1’s preferential recognition of H2A.Z, we mapped the binding interface using CSP of SUV420H1–NCP^167^ complexes. Based on backbone assignments for H2A and H2A.Z in complex with SUV420H1 (Figure S7A, B), we calculated CSP to obtain site-specific interaction information (Figure 5A). The CSP analysis not only confirmed and refined known interaction sites within the acidic patch but also identified two additional perturbed regions. One was the QF motif, which is conserved in both H2A.Z and H2A. The other was the M2 region, which exhibited pronounced CSP specifically in H2A.Z. This finding suggests that the M2 region contributes to SUV420H1 recognition in an H2A.Z-selective manner.

**Figure 5.**
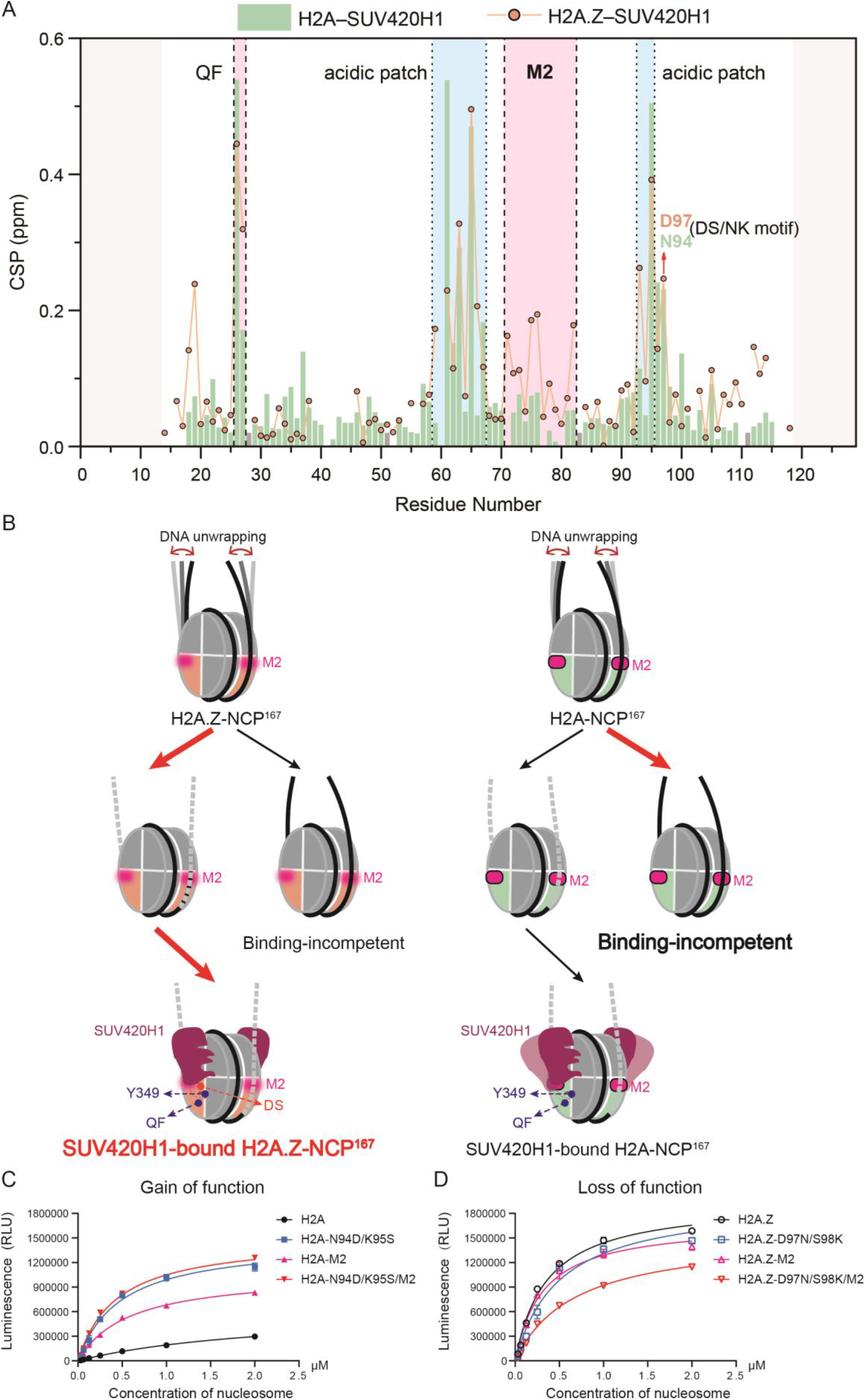
Dynamic differences in the M2 region underlie SUV420H1’s preferential recognition of H2A.Z. (A) Chemical shift perturbation (CSP) analysis of SUV420H1 binding to H2A.Z and H2A nucleosomes. In addition to validating and refining the known acidic patch interactions, solid-state NMR identified two previously unreported perturbed regions: the QF motif (present in both H2A and H2A.Z) and the M2 region (exhibiting pronounced CSP exclusively in H2A.Z). (B) Working model in which M2 dynamics and the DS motif mediate H2A.Z-selective recognition, whereas the conserved Y349–QF contact supports catalytic engagement. SUV420H1 senses the M2-facilitated partial DNA unwrapping of H2A.Z nucleosomes, which underlies variant-selective recognition. (C) The MTase-Glo analysis results of the HMT assay carried out using wild-type SUV420H1 on nucleosomes containing H2A, H2A-N94D/K95S, H2A-M2, or H2A-N94D/K95S/M2. Introducing the H2A.Z M2 sequence into H2A (H2A-M2) significantly enhances SUV420H1 catalytic activity, indicating that the H2A.Z-derived M2 sequence is sufficient to confer partial H2A.Z-like activity into H2A. The experiments were replicated three times, and the error bars were calculated as the standard error. (D) The MTase-Glo analysis results of the HMT assay carried out using wild-type SUV420H1 on nucleosomes containing H2A.Z, H2A.Z-D97N/S98K, H2A.Z-M2, or H2A.Z-D97N/S98K/M2. Replacing the H2A.Z M2 region with the H2A counterpart (H2A.Z-M2) reduces SUV420H1 catalytic activity. The experiments were replicated three times, and the error bars were calculated as the standard error. The reciprocal DS/NK swaps combined with the M2 swaps (C and D) reveal that the M2 and DS regions exert additive effects on activity, suggesting that both regions synergistically contribute to SUV420H1’s preferential recognition of H2A.Z.

In the reported structures of the SUV420H1–H2A.Z and SUV420H1–H2A nucleosome complexes, the M2 region does not directly contact the enzyme^55,56^. Instead, these structures show pronounced nucleosomal DNA unwrapping upon SUV420H1 binding^55,56^. Given our demonstration that the intrinsic flexibility of the M2 region promotes DNA-end accessibility, we propose that the H2A.Z-specific CSP observed in the M2 region reflects SUV420H1 sensing the pre-existing conformational landscape of H2A.Z nucleosomes rather than engaging M2 as a simple static binding site. In this model, the M2 dynamic element predisposes H2A.Z nucleosomes to partially unwrapped conformations, thereby lowering the energetic barrier for the DNA deformation required for SUV420H1 binding and catalysis (Figure 5B).

To assess the functional significance of the M2 region, we performed SUV420H1 enzymatic assays with different chimeric H2A/H2A.Z nucleosomes. Simply swapping the previously highlighted DS motif in H2A.Z with the corresponding NK motif in H2A did not fully reverse SUV420H1’s substrate preference (Figure S8), suggesting that additional factors beyond the DS/NK motif contribute to recognition specificity. Strikingly, replacing the entire M2 region of H2A with that of H2A.Z (H2A-M2) significantly enhanced SUV420H1 catalytic activity (Figure 5C), whereas introducing the H2A counterpart region into H2A.Z (H2A.Z-M2) reduced activity (Figure 5D). Moreover, combining M2 exchange with reciprocal DS/NK substitutions produced stronger effects than either alteration alone (Figure 5C, D). Thus, the M2 region and the extended-acidic-patch-associated DS motif act together to promote SUV420H1’s preferential recognition and catalytic activity toward H2A.Z nucleosomes (Figure 5B).

Separately, CSP-guided mutagenesis identified a conserved QF-associated contact involving SUV420H1 Y349 that supports catalytic activity but does not account for H2A.Z selectivity, further distinguishing general catalytic engagement from M2-dependent dynamic recognition (Figure 5B; Figure S7C; Figure S9).

## Discussion

H2A.Z and H2A mediate distinct chromatin functions that cannot be explained by static structures alone. Using ^1^H-detected ssNMR under fast MAS, we resolved that H2A.Z and H2A, despite their near-identical folds, exhibit markedly divergent backbone dynamics, specifically enhanced flexibility in the L1 loop (M1, R39– V46) and the α2–L2 region (M2, N71–V78). Chimeric segment-swapping established that local amino acid sequences encode the flexibility of both regions. We further demonstrate that the inherent mobility of the M2 region drives nucleosomal DNA unwrapping and determines SUV420H1’s variant-selective recognition, identifying sequence-encoded dynamics as the physical basis for this epigenetic preference.

Structural studies have identified subtle architectural differences between H2A.Z- and H2A-containing nucleosomes, including the L1 loop, acidic patch, and docking domain^29,34,35,37^, yet whether H2A.Z stabilizes or destabilizes the nucleosome has remained controversial^16,29–34,36,38^. This persistent ambiguity suggests that static architectural comparison alone does not fully account for their distinct functional behaviors. Our ssNMR measurements provide direct experimental evidence that H2A.Z exhibits enhanced backbone flexibility in the L1 loop (M1) and the α2–L2 region (M2), establishing sequence-encoded dynamics as a determinant of variant-specific function.

Previous MNase assays demonstrated increased DNA accessibility in H2A.Z nucleosomes^36^, but the determinants within H2A.Z responsible for this effect remained elusive. By integrating MNase assays with M2-swap mutants, we show that M2 region flexibility is a major driver of enhanced DNA unwrapping. Importantly, M2 dynamics persist whether DNA ends are stabilized by linker histone H1 or detached by SUV420H1^55,56,62–67^, indicating that M2 flexibility actively contributes to DNA accessibility rather than passively reflecting DNA end conformation. This provides experimental support for the view that H2A.Z incorporation lowers the energetic barrier for DNA breathing^38^and establishes a residue-level direct link between backbone dynamics and nucleosome accessibility.

For SUV420H1 recognition, our NMR data reveal that the dynamic M2 region serves as a previously unrecognized determinant of variant-selective binding, extending beyond models that attribute specificity solely to the DS motif^22,55^. We show that SUV420H1 reads the H2A.Z-specific conformational landscape shaped by M2 backbone flexibility to distinguish otherwise architecturally similar nucleosomes. Additionally, ssNMR uncovers a functionally essential Y349–QF contact that appears too distal for direct engagement in cryo-EM structures, revealing a hidden interaction required for catalytic activity. These findings establish that local nucleosome dynamics can function as a specificity filter, adding a distinct layer of epigenetic control beyond static sequence determinants.

Collectively, our findings define a sequence-dynamics-function axis that translates local histone sequence into variant-selective chromatin behavior. By establishing that the intrinsic flexibility of the H2A.Z M2 region actively controls DNA accessibility and enzyme recognition, we demonstrate that conformational dynamics themselves constitute regulatory determinants operating alongside covalent marks and static structure. The resulting increase in nucleosomal DNA breathing may facilitate both epigenetic enzyme engagement and transcription machinery access^68^, accounting for the characteristic enrichment of H2A.Z at genomic sites requiring elevated DNA accessibility^17,19,20,23^. We propose that such dynamical encoding represents a general principle by which histone variants and chromatin factors diversify their functional repertoire.

The present work defines a mechanistic baseline in a precisely controlled mononucleosome system, a reductionist setting chosen to isolate the intrinsic sequence-dynamics relationship of the H2A.Z M2 region. This controlled foundation now positions us to systematically integrate the additional layers of native chromatin complexity—including histone post-translational modifications, variable linker DNA lengths, neighboring nucleosome contacts, and the competitive binding of chromatin-associated factors. Extending the ssNMR platform to nucleosome arrays will be particularly important for testing whether M2-driven DNA unwrapping is modulated by higher-order compaction. Likewise, applying this dynamic mapping strategy to other histone variants promises to reveal whether sequence-encoded flexibility represents a general regulatory vocabulary in chromatin biology.

## Acknowledgment

This work was supported by grants from the National Key R&D Program (2025YFA1308801) and the National Natural Science Foundation of China (31971128, 32400459, 32320103008, 32270651, 32550379), CAS Project for Young Scientists in Basic Research, Grant No. YSBR-068. CAS Strategic Priority Research Program (XDB1000000).

## Author Contributions Statement

T.Z., L.H., Z.Z., and S.X. conceived and designed the experiments. T.Z. performed most of the biochemical and NMR experiments of NCP. L.H. purified SUV420H1 and performed the biochemical experiments related to SUV420H1. X.L. processed the relaxation data. B.L assisted with the preparation of NCP samples. J.L. and C.S. helped with the NMR experiments. S.F. assisted with the preparation of histone proteins samples. T.Z., L.H., Z.Z., and S.X. wrote the manuscript.

