## Supporting information for "Sequence-encoded H2A.Z nucleosome dynamics control DNA unwrapping and SUV420H1 recognition"

Table of contents

1: Method

1.1: Mononucleosome sample production

1.2: Assembly of nucleosome complexes

1.3: Sedimentation of nucleosomes for solid-state NMR

1.4: The MTase-Glo methyltransferase assay of SUV420H1

1.5: Micrococcal nuclease (MNase) treatment assay

1.6: Solid-state NMR spectroscopy

1.7: Backbone resonance assignment

1.8: Data analysis and error estimation of $R_{1\rho}$

1.9: Chemical shift perturbation analysis

Table S1. Dipolar coupling parameters from $^{1}$H-detected solid-state NMR for the H2A.Z sample series.

Table S2. Dipolar coupling parameters from $^{1}$H-detected solid-state NMR for the H2A sample series.

Table S3. $^{1}$H-detected solid-state NMR spectral width of the H2A.Z sample series.

Table S4. $^{1}$H-detected solid-state NMR spectral width of the H2A sample series.

Table S5. $^{1}$H-detected solid-state NMR acquisition time of the H2A.Z sample series.

Table S6. $^{1}$H-detected solid-state NMR acquisition time of the H2A sample series.

2: Supplementary results

S1. Preparation and characterization of nucleosome samples.

S2. Backbone resonance assignment and secondary structure analysis of H2A.Z.

S3. Backbone resonance assignment and secondary structure analysis of H2A.Z chimeras.

S4. Backbone resonance assignment and secondary structure analysis of H2A.

S5. Backbone resonance assignment and secondary structure analysis of the H2A-M2 chimera.

S6. H1.4 does not perturb the core structural domains of H2A or H2A.Z.

S7. Backbone resonance assignments of H2A and H2A.Z in SUV420H1–NCP$^{167}$ complexes.

S8. Bidirectional exchange of the DS and NK motifs did not fully reverse SUV420H1 activity levels between H2A.Z and H2A.

S9. Residue Y349 in SUV420H1 mediates nucleosome binding and catalytic activity.

3: References

### 1: Method

#### 1.1 Mononucleosome sample preparation

Mononucleosomes were prepared following established protocols adapted from Dyer et al[1]. The detailed procedures are described below.

##### Histone purification

Recombinant human histones were expressed in *E. coli* BL21 (DE3) cells. H2A, H2B, and H3 were cloned into the pET-30a vector, while H2A.Z and H4 were cloned into the pET-3a vector. Bacterial cultures were grown to an $OD_{600}$ of 0.6–0.8, and protein expression was induced with 0.5 mM isopropyl β-D-1-thiogalactopyranoside (IPTG) at 37 °C for 6 h. Cells were harvested by low-speed centrifugation (4,300 × *g*) and lysed using a low-temperature, high-pressure cell disruptor. The target proteins were predominantly found in insoluble inclusion bodies. Following high-speed centrifugation (19,000 × *g*), the inclusion bodies were solubilized in unfolding buffer (50 mM $Na_2HPO_4$, pH 7.2, 6 M guanidine hydrochloride) with further sonication to ensure complete dissolution. The solubilized sample was clarified by another high-speed centrifugation step (19,000 × *g*). The clear supernatant was then dialyzed overnight against a denaturing buffer (50 mM $Na_2HPO_4$, pH 7.2, 8 M urea, 100 mM NaCl) at 4 °C. Subsequently, histones were sequentially purified by cation-exchange chromatography using a gravity column packed with SP resin (BBI) and reversed-phase high-performance liquid chromatography (HPLC). Purified histones were lyophilized to powder and stored at -80 °C until use. Histone chimeric mutants (H2A.Z-M1, H2A.Z-M2, H2A-M2) were prepared using identical procedures. For $^{1}H$-detected NMR studies, isotopically labeled histones were expressed in $D_2O$-based LR medium (100% $D_2O$, 2.5 g/L $^{1}H$-$^{13}C$ glucose, 0.5 g/L $^{15}NH_4Cl$). Unlabeled histones were produced in LB medium.

##### DNA preparation

The pWM530 plasmid, containing 12 tandem repeats of the 167 bp Widom 601 sequence[2], was transformed into *E. coli* DH5α and amplified by large-scale culture in rich TB medium. Plasmid DNA was extracted using the alkaline lysis method, followed by purification steps including isopropanol precipitation, RNase A (QIAGEN) digestion, PEG 6000 precipitation, phenol-chloroform extraction, and absolute ethanol precipitation. The purified plasmid was digested with ScaI restriction endonuclease (Thermo Fisher Scientific). The digestion product was further purified by PEG 6000 precipitation, absolute ethanol precipitation, and anion-exchange chromatography on a 5 mL HiTrap Capto Q column (GE Healthcare). The resulting high-quality 167 bp DNA fragment was dissolved in TE 10/0.1 buffer (10 mM Tris-HCl, pH 7.5, 0.1 mM EDTA) and used for subsequent nucleosome assembly.

##### Histone octamer preparation

Lyophilized histone powders (H2A or H2A.Z, H2B, H3, H4) were each dissolved in unfolding buffer, mixed in equimolar ratios, and diluted to a final concentration of approximately 1 mg/mL. The mixture was refolded by dialysis against refolding buffer (10 mM Tris-HCl, pH 7.5, 2 M KCl, 1 mM EDTA, 5 mM DTT) at 4°C, allowing proper folding and octamer assembly. After dialysis, the assembly product was purified by size-exclusion chromatography on a Superdex 200 Increase column (GE Healthcare) to obtain histone octamers. Octamer assembly efficiency was assessed by 18% SDS-PAGE.

##### Mononucleosome reconstitution

Mononucleosomes were assembled using the salt gradient dialysis method. First, the optimal molar ratio of histone octamer to 167 bp DNA (typically between 0.8:1 and 1.4:1) was determined through small-scale pilot experiments.

Large-scale reconstitution was then performed using this optimal ratio. The mixture of histone octamer and DNA fragment was placed into a dialysis bag and dialyzed against high-salt refolding buffer at 4°C. Concurrently, a peristaltic pump was used to continuously and slowly introduce low-salt TE 10/1 buffer (10 mM Tris-HCl, pH 7.5, 1 mM EDTA, 5 mM DTT) into the refolding buffer, gradually reducing the KCl concentration from 2 M to 0.6 M over approximately 17–18 h. Finally, the sample was transferred to HE buffer (10 mM HEPES, pH 7.5, 0.1 mM EDTA, 5 mM DTT) and dialyzed for an additional 5 h. Assembled mononucleosomes were analyzed by 5% native PAGE.

**1.2 Preparation of Nucleosome Complexes**

**Purification of linker histone H1.4**

Human H1.4 was expressed in *E. coli* BL21 (DE3) using LB medium. Cultures were grown at 37°C to an $OD_{600}$ of 0.6–0.8, and expression was induced with 0.5 mM IPTG for 6 h. After cell lysis by high-pressure, the SUMO-tagged H1.4 was initially purified by $Ni^{2+}$ affinity chromatography (Union Biotech). The SUMO tag was cleaved by ULP1 protease, and highly pure H1.4 was obtained by a subsequent cation-exchange chromatography step on a 1 mL HiTrap SP HP column (GE Healthcare). The purified H1.4 was dialyzed into storage buffer (20 mM Tris-HCl, pH 7.5, 100 mM NaCl, 2 mM EDTA, 20% glycerol), aliquoted, and stored at -80°C.

**Preparation of H1.4–nucleosome complex**

The H1.4–nucleosome complex (H1.4–$NCP^{167}$) was prepared by incorporating H1.4 during the mononucleosome reconstitution process. Specifically, when the KCl concentration reached 0.6 M during the salt gradient dialysis, purified H1.4 protein was added to the nucleosome solution at an optimized molar ratio. Dialysis was continued for another 3 h, followed by a final dialysis step against HE buffer for 5 h to obtain the H1.4-$NCP^{167}$ complex.

**Purification of SUV420H1**

SUV420H1 was purified following a previously described protocol[3]. DNA sequence encoding human SUV420H1 (aa: 1–393) with a C-terminal 6×His tag was amplified by PCR and subcloned into *E. coli* expression vector pET17b. The resulting construct was transformed into BL21 Rosetta (DE3) for recombinant protein expression. Transformed cells were grown in LB medium at 37 °C until reaching an $OD_{600}$ of nearly 0.8, followed by induction with 0.2 mM IPTG and incubation at 16 °C overnight. Cells were harvested by centrifugation and resuspended in SUV420H1 lysis buffer (20 mM MES pH 6.5, 500 mM NaCl, 5% glycerol, 20 mM imidazole, 5 mM β-mercaptoethanol). Following cell disruption and clarification, the lysate was incubated with Ni-NTA agarose resin (QIAGEN) for 1 h at 4°C. The resin was subsequently transferred to a gravity-flow column and washed with 150 mL lysis buffer to remove nonspecifically bound proteins. Bound SUV420H1 was eluted using SUV420H1 elution buffer (20 mM MES, pH 6.5, 500 mM NaCl, 5% glycerol, and 250 mM imidazole, 5 mM β-mercaptoethanol).

The eluate was diluted to reduce the NaCl concentration to 200 mM and loaded onto a 5 mL HiTrap SP HP cation-exchange column (GE Healthcare) pre-equilibrated with SUV420H1 buffer A (20 mM MES, pH 6.5, 200 mM NaCl, 5% glycerol, and 5 mM β-mercaptoethanol). SUV420H1 was eluted using a 0.2–1 M linear NaCl gradient. Fractions containing SUV420H1 were pooled and further purified by size-exclusion chromatography on a Superdex 200 column (GE Healthcare) equilibrated with SUV420H1 Superdex buffer (20 mM MES, pH 6.5, 300 mM NaCl, 5% glycerol, and 2 mM DTT). Purified SUV420H1 was concentrated as required, flash-frozen in liquid nitrogen, and stored at -80°C until further use.

**Preparation of SUV420H1–nucleosome complex**

The SUV420H1–nucleosome complex (SUV420H1–$NCP^{167}$) was prepared by mixing mononucleosomes and purified SUV420H1 at a molar ratio of 1:2.2. Assembled mononucleosomes were diluted to 0.05 mg/mL (DNA concentration) with HE buffer containing 5% glycerol. Purified SUV420H1 protein was then added slowly to achieve a final molar ratio of 2.2:1 (SUV420H1: nucleosome). After thorough mixing, the sample was immediately dialyzed against HE buffer containing 5% glycerol at 4°C for 3 h. Following dialysis, the complex sample was centrifuged (20,000 × *g*) to remove any precipitate. The supernatant was collected and concentrated, then subjected to a second high-speed centrifugation (20,000 × *g*) to remove any precipitate formed during concentration. The final clear supernatant, containing SUV420H1–NCP$^{167}$, was used for subsequent experiments. A protease inhibitor cocktail (Roche) was added to the complex sample to ensure stability.

##### 1.3 Sedimentation of nucleosomes for solid-state NMR

We utilized an in-house developed ultracentrifugation filling tool, compatible with various Bruker solid-state NMR rotors, for sedimentation NMR studies[4,5]. This tool withstands prolonged ultracentrifugation and was employed to sediment nucleosomes directly into 1.3 mm rotors. Approximately 2 mg of the solution-state nucleosome sample was transferred into the funnel-shaped inner chamber of the filling tool for the 1.3 mm rotor. After precise balancing, the assembly was subjected to ultracentrifugation at 240,000 × *g* for 60 h at 4°C using a swinging bucket rotor (MLS-50), sedimenting the nucleosomes into the solid-state NMR rotor. Following centrifugation, the supernatant was carefully removed, and the rotor containing the sedimented, labeled nucleosomes was retrieved. After capping with the turbine lid, the rotor was ready for solid-state NMR experiments. Sedimentation efficiency was assessed by calculating the ratio of DNA concentration in the supernatant post-centrifugation to that in the initial sample, which consistently exceeded 95%.

##### 1.4 The MTase-Glo methyltransferase assay of SUV420H1

The methyltransferase activity of SUV420H1 was quantified using the MTase-Glo™ Methyltransferase Assay (Promega, Cat# V7601). Reactions containing 150 nM SUV420H1 and increasing concentrations of nucleosome substrates (0–2 μM) were incubated in 50 mM Tris-HCl (pH 7.5) at 30°C for 60 min. Reactions were quenched by adding TFA to a final concentration of 0.1%. The resulting SAH was converted to ATP through the MTase-Glo detection system, and ATP production was quantified by a luciferase-coupled luminescence assay using an EnSight Multimode Plate Reader (PerkinElmer). All experiments were independently repeated three times, and data are presented as mean ± SD. Apparent kinetic parameters were obtained by fitting the data to the Michaelis-Menten equation using GraphPad Prism 9.0. Differences between fitted curves were assessed using the curve comparison function implemented in GraphPad Prism 9.0.

##### 1.5 Micrococcal Nuclease (MNase) treatment assay

MNase treatment assays were performed using nucleosome core particles reconstituted with 147 bp DNA, prepared using the same method as for the mononucleosomes assembled on 167 bp DNA described above. Nucleosomes (1 μg, quantified by DNA) were incubated with 5 gel units of MNase (Beyotime, Cat# D7201S) at 37 °C in a 50 μl reaction volume. Different incubation times were used based on experimental requirements. For H2A.Z, H2A.Z-M2, H2A, and H2A-M2, time points were 0 min and 50 min. For H2A.Z, H2A.Z–SUV420H1, H2A, and H2A–SUV420H1, time points were 0 min and 30 min. At each designated time point, a 20 μL aliquot was mixed with 10 μL of reaction termination buffer and incubated overnight at 65 °C. The reaction termination buffer contained 10 mM HEPES (pH 7.5), 71.45 mM EDTA, 5 mM DTT, 0.1% SDS, and 0.5 mg/mL proteinase K. Finally, reaction products were analyzed by electrophoresis on 10% native PAGE (Sangon Biotech, Cat# B650003-0001) in 1×TBE buffer. DNA bands were visualized by staining with 4S Red Plus nucleic acid stain (Sangon Biotech, Cat# A606695), and quantitative analysis was performed using ImageJ software. All experiments were independently repeated three times, and data are presented as mean + SD.

**1.6 Solid-state NMR spectroscopy**

All $^{1}H$-detected solid-state NMR experiments were performed on a Bruker wide-bore NEO spectrometer operating at a $^{1}H$ frequency of 600 MHz, equipped with a 1.3 mm HXY triple-resonance magic-angle spinning (MAS) probe. Samples were spun at a MAS frequency of 60 kHz, and the effective sample temperature was maintained at approximately 22°C. A suite of multi-dimensional cross-polarization-based (CP-based) spectra was acquired for backbone resonance assignment, including 2D NH correlation spectra and 3D CANH, CA(CO)NH, CONH, CO(CA)NH, and CBCANH spectra. Detailed acquisition parameters for each spectrum are provided in the supporting information (Tables S1–S6). The cross-polarization (CP) power transmitted from Cα to Cβ is 30 kHz. 3D experiments were recorded using a block acquisition strategy, with each 3D dataset collected in a block of approximately one day. The $^{1}H$ signal of a small molecule at around 2.8 ppm was measured between blocks to monitor and correct for external magnetic field drift. $^{15}N$ transverse relaxation rates ($R_{1\rho}$) were measured using a modified 3D experiment. A variable delay list (vdlist) was incorporated into a standard CP-based 2D NH correlation pulse sequence, effectively converting it into a 3D experiment. A $^{15}N$ spin-lock field strength of 9 kHz was applied for durations of 0.02, 10, 20, 40, and 60 ms. $R_{1\rho}$ values were extracted by fitting the decay of signal intensity as a function of the spin-lock time. All solid-state NMR spectra were acquired using TopSpin 4.1.4 software. Data were processed with NMRPipe[6] and analyzed using Sparky[7]. To enhance spectral resolution, the number of points in both indirect dimensions was doubled using the smile script during processing.

**1.7 Backbone resonance assignment**

Backbone resonance assignments for human H2A.Z (pH 6.5) were performed manually using standard sequential assignment methodologies. Assignments were primarily based on linking inter-residue and intra-residue correlations provided by Cα and CO information in 3D CANH, CA(CO)NH, CONH, and CO(CA)NH spectra. Cβ information from a 3D CBCANH spectrum employing the $POST - C4_{16}^{1}$ (PC4) pulse sequence was used to aid amino acid type identification and validate assignment correctness. PC4 was originally designed for moderate MAS rates (10–20 kHz) to enhance Cα–Cβ magnetization transfer while suppressing nontarget signals[8]. Here, we extended PC4 to 60 kHz MAS in a 3D CBCANH experiment. Compared to the conventional DREAM sequence, the PC4-based CBCANH provided more reliable Cβ information for a greater number of residues, significantly enhancing the reliability and completeness of the H2A.Z backbone assignment. Ultimately, 87 residues within the rigid region (14–118) of H2A.Z were unambiguously assigned in the CP-based spectra, covering 85% of the 102 assignable non-proline residues in this region.

Given the high sequence homology between human H2A and Drosophila melanogaster H2A, assignments for human H2A were transferred using previously published NMR data for Drosophila H2A[9]. CO information from 3D CONH spectra was used for assignment transfer. To ensure assignment accuracy, this was corroborated by sequential linking using Cα correlations from CANH (intra-residue) and CA(CO)NH (preceding residue). Using this approach, 92 out of 94 non-proline residues (98%) in the rigid region (16–113) of human H2A were assigned. Backbone assignments for other solid-state NMR samples, including H2A.Z (pH 7.5), H2A.Z-M1, H2A.Z-M2, H2A-M2, SUV420H1-bound H2A.Z, and SUV420H1-bound H2A, were achieved by transfer from 3D CANH spectra. Secondary structure analysis was performed using the assigned chemical shifts with the δ2D web server （https://www-cohsoftware.ch.cam.ac.uk/index.php/d2D）.

The signal intensity ratio in the CANH spectra was calculated as $I_b/I_0$, where $I_b$ corresponds to the peak intensity of each residue, and $I_0$ corresponds to that of a reference residue, Q26 in H2A.Z (or Q24 in H2A). The error associated with $I_b/I_0$ was calculated according to Equation (1):

$$\sigma = \frac{I_b}{I_0} \times \sqrt{\left(\frac{1}{SNR_b}\right)^2 + \left(\frac{1}{SNR_0}\right)^2} \tag{1}$$

where $SNR_b$ and $SNR_0$ denote the signal-to-noise ratios of each residue and the reference residue, respectively.

##### 1.8 Data analysis and error estimation of $R_{1\rho}$

$^{15}$N $R_{1\rho}$ relaxation rates were determined by fitting the exponential decay curve of the signal intensity as a function of relaxation time. The relationship between signal intensity and relaxation time is described by Equation (2):

$$I(t) = I_0 \cdot exp\left(-R_{1\rho} \cdot t\right) \tag{2}$$

where $I(t)$ is the signal intensity measured at spin-lock time t, $I_0$ is the initial signal intensity at t = 0, and $R_{1\rho}$ is the relaxation rate (in $s^{-1}$).

The error in the decay of the normalized signal intensity for the representative residues Q26 and L76 in H2A.Z was determined by accounting for the experimental noise level. To accurately assess the uncertainty in the extracted $R_{1\rho}$ values, Monte Carlo simulations were performed. The experimental noise level was quantified by measuring the root-mean-square (rms) noise in a signal-free region of the spectrum using Sparky. For each relaxation decay curve, 100,000 synthetic datasets were generated by adding random noise, matching the experimental rms noise level, to the original peak intensities. Each synthetic dataset was fitted using non-linear least squares (scipy.optimize.curve_fit in Python). The final reported $R_{1\rho}$ value and its error are the mean and standard deviation, respectively, of the distribution of the 100,000 fitted rates. This method provides a robust assessment of parameter precision by incorporating the actual noise characteristics of the experimental data.

##### 1.9 Chemical shift perturbation analysis

Protein–nucleosome interactions were characterized by quantifying chemical shift perturbation (CSP). In this study, CSP was calculated according to Equation (3):

$$CSP = \sqrt{(\Delta\delta_H)^2 + (\Delta\delta_N/5.4)^2} \tag{3}$$

where $\Delta\delta_H$ and $\Delta\delta_N$ are the chemical shift changes (in ppm) observed in the $^1$H and $^{15}$N dimensions, respectively. A scaling factor of 5.4 was applied to the $^{15}$N chemical shift changes to account for the difference in chemical shift dispersion between $^1$H and $^{15}$N nuclei.

**Table S1. Dipolar coupling parameters from $^1$H-detected solid-state NMR for the H2A.Z sample series.**

| | **Transfer steps** | **H→N** | **N→H$^N$** | **H→Cα** | **Cα→N** | **H→CO** | **CO→N** | **Cα→CO** | **CO→Cα** |
|---|---|---|---|---|---|---|---|---|---|
| | **Ramp** | 80–100 % ramp on H | 100–80 % ramp on H | 90–100 % ramp on H | 90–100 % ramp on Cα | 90–100 % ramp on H | 90–100 % ramp on CO | 90–100 % ramp on CO | 90–100 % ramp on Cα |
| **H2A.Z (pH6.5)** | CP power (kHz) | H:116.16<br>N:41.82 | N:41.82<br>H:116.16 | H:105.41<br>Cα:45.64 | Cα:40.25<br>N:20.75 | H:99.60<br>CO:40.25 | CO:40.25<br>N:20.75 | 29.46 | 29.46 |
| | Contact time(us) | 500 | 500 | 4000 | 6000 | 3500 | 6000 | 3500 | 4500 |
| **H2A.Z (pH7.5)** | CP power (kHz) | H:111.89<br>N:42.79 | N:42.79<br>H:112.68 | H:102.72<br>Cα:45.64 | Cα:40.25<br>N:21.23 | | | | |
| | Contact time(us) | 1000 | 500 | 4000 | 6000 | | | | |
| **H2A.Z-M1** | CP power (kHz) | H:115.88<br>N:43.81 | N:43.81<br>H:116.63 | H:108.90<br>Cα:45.64 | Cα:40.25<br>N:21.04 | | | | |
| | Contact time(us) | 1500 | 500 | 4000 | 6000 | | | | |
| **H2A.Z-M2** | CP power (kHz) | H:110.67<br>N:43.81 | N:43.81<br>H:109.12 | H:106.76<br>Cα:45.64 | Cα:40.25<br>N:22.40 | H:98.47<br>CO:40.25 | CO:40.25<br>N:22.40 | | |
| | Contact time(us) | 1000 | 500 | 4500 | 6000 | 5000 | 6000 | | |
| **H2A.Z–H1** | CP power (kHz) | H:116.16<br>N:43.30 | N:43.30<br>H:114.44 | | | | | | |
| | Contact time(us) | 1500 | 500 | | | | | | |
| **H2A.Z–SUV420H1** | CP power (kHz) | H:117.95<br>N:42.79 | N:42.79<br>H:114.62 | H:104.94<br>Cα:45.64 | Cα:40.25<br>N:20.56 | | | | |
| | Contact time(us) | 1000 | 500 | 4500 | 6000 | | | | |
| **H2A.Z–SUV420H1-Y349A** | CP power (kHz) | H:116.16<br>N:43.81 | N:43.81<br>H:116.16 | | | | | | |
| | Contact time(us) | 1250 | 500 | | | | | | |

**Table S2. Dipolar coupling parameters from $^{1}$H-detected solid-state NMR for the H2A sample series.**

| | **Transfer steps** | **H→N** | **N→H$^{N}$** | **H→Cα** | **Cα→N** | **H→CO** | **CO→N** | **Cα→CO** |
|---|---|---|---|---|---|---|---|---|
| | **Ramp** | 80–100 % ramp on H | 100–80 % ramp on H | 90–100 % ramp on H | 90–100 % ramp on Cα | 90–100 % ramp on H | 90–100 % ramp on CO | 90–100 % ramp on CO |
| **H2A** | CP power (kHz) | H:114.99 N:43.29 | N:43.29 H:111.79 | H:104.25 Cα:45.64 | Cα:45.64 N:20.80 | H:97.98 CO:40.25 | CO:38.03 N:23.41 | 28.46 |
| | Contact time(us) | 1500 | 500 | 5000 | 6000 | 4000 | 6000 | 4500 |
| **H2A-M2** | CP power (kHz) | H:118.05 N:43.81 | N:43.81 H:116.47 | H:106.47 Cα:45.64 | Cα:39.53 N:21.04 | | | |
| | Contact time(us) | 1500 | 500 | 3500 | 5500 | | | |
| **H2A–H1** | CP power (kHz) | H:120.29 N:43.30 | N:43.30 H:116.16 | | | | | |
| | Contact time(us) | 1500 | 500 | | | | | |
| **H2A – SUV420H1** | CP power (kHz) | H:115.77 N:43.81 | N:43.81 H:110.98 | H:104.25 Cα:45.64 | Cα:38.79 N:22.40 | | | |
| | Contact time(us) | 2000 | 500 | 4000 | 6000 | | | |

**Table S3. $^{1}$H-detected solid-state NMR spectral width of the H2A.Z sample series**.

| Spectrum | Nucleus | Spectral width (ppm) | | | | | | |
|---|---|---|---|---|---|---|---|---|
| | | H2A.Z (pH6.5) | H2A.Z (pH7.5) | H2A.Z-M1 | H2A.Z-M2 | H2A.Z–H1 | H2A.Z – SUV420H1 | H2A.Z – SUV420H1 -Y349A |
| 2D NH | N | 32 | 32 | 32 | 32 | 32 | 32 | 32 |
| | $H^N$ | 19.8368 | 19.8368 | 19.8368 | 19.8368 | 19.8368 | 19.8368 | 19.8368 |
| 3D CANH | Cα | 26 | 26 | 26 | 26 | | 26 | |
| | N | 32 | 32 | 32 | 32 | | 32 | |
| | $H^N$ | 19.8368 | 19.8368 | 19.8368 | 19.8368 | | 19.8368 | |
| 3D CONH | CO | 11 | | | 11 | | | |
| | N | 32 | | | 32 | | | |
| | $H^N$ | 19.8368 | | | 19.8368 | | | |
| 3D CA(CO)NH | Cα | 26 | | | | | | |
| | N | 32 | | | | | | |
| | $H^N$ | 19.8368 | | | | | | |
| 3D CO(CA)NH | CO | 11 | | | | | | |
| | N | 32 | | | | | | |
| | $H^N$ | 19.8368 | | | | | | |
| 3D CBCANH | C | 62 | | | | | | |
| | N | 32 | | | | | | |
| | $H^N$ | 19.8368 | | | | | | |

**Table S4. $^{1}$H-detected solid-state NMR spectral width of the H2A sample series.**

| Spectrum | Nucleus | Spectral width (ppm) | | | |
|---|---|---|---|---|---|
| | | H2A | H2A-M2 | H2A–H1 | H2A–SUV420H1 |
| 2D NH | N | 32 | 32 | 32 | 32 |
| | $H^N$ | 19.8368 | 19.8368 | 19.8368 | 19.8368 |
| 3D CANH | Cα | 26 | 26 | | 26 |
| | N | 32 | 32 | | 32 |
| | $H^N$ | 19.8368 | 19.8368 | | 19.8368 |
| 3D CONH | CO | 11 | | | |
| | N | 32 | | | |
| | $H^N$ | 19.8368 | | | |
| 3D CA(CO)NH | Cα | 26 | | | |
| | N | 32 | | | |
| | $H^N$ | 19.8368 | | | |

**Table S5. [1]H-detected solid-state NMR acquisition time of the H2A.Z sample series.**

| Spectrum | Nucleus | Acquisition time (ms) | | | | | | |
|---|---|---|---|---|---|---|---|---|
| | | H2A.Z (pH6.5) | H2A.Z (pH7.5) | H2A.Z-M1 | H2A.Z-M2 | H2A.Z–H1 | H2A.Z – SUV420H1 | H2A.Z – SUV420H1 -Y349A |
| 2D NH | N | 29.8 | 29.8 | 29.8 | 29.8 | 29.8 | 29.8 | 29.8 |
| | $H^N$ | 20.0 | 20.0 | 20.0 | 20.0 | 20.0 | 20.0 | 20.0 |
| 3D CANH | Cα | 4.6 | 4.6 | 4.6 | 4.6 | | 4.6 | |
| | N | 8.7 | 8.7 | 8.7 | 8.7 | | 8.7 | |
| | $H^N$ | 15.0 | 15.0 | 15.0 | 15.0 | | 15.0 | |
| 3D CONH | CO | 6.0 | | | 6.0 | | | |
| | N | 8.7 | | | 8.7 | | | |
| | $H^N$ | 15.0 | | | 15.0 | | | |
| 3D CA(CO)NH | Cα | 4.6 | | | | | | |
| | N | 8.7 | | | | | | |
| | $H^N$ | 15.0 | | | | | | |
| 3D CO(CA)NH | CO | 6.0 | | | | | | |
| | N | 8.7 | | | | | | |
| | $H^N$ | 15.0 | | | | | | |
| 3D CBCANH | C | 2.1 | | | | | | |
| | N | 7.7 | | | | | | |
| | $H^N$ | 15.0 | | | | | | |

**Table S6. $^{1}$H-detected solid-state NMR acquisition time of the H2A sample series.**

| Spectrum | Nucleus | Acquisition time (ms) | | | |
|---|---|---|---|---|---|
| | | H2A | H2A-M2 | H2A–H1 | H2A–SUV420H1 |
| 2D NH | N | 29.8 | 29.8 | 29.8 | 29.8 |
| | $H^N$ | 20.0 | 20.0 | 20.0 | 20.0 |
| 3D CANH | Cα | 4.6 | 4.6 | | 4.6 |
| | N | 8.7 | 8.7 | | 8.7 |
| | $H^N$ | 15.0 | 15.0 | | 15.0 |
| 3D CONH | CO | 6.0 | | | |
| | N | 8.7 | | | |
| | $H^N$ | 15.0 | | | |
| 3D CA(CO)NH | Cα | 4.6 | | | |
| | N | 8.7 | | | |
| | $H^N$ | 15.0 | | | |

#### 2: Supplementary Figure Legends

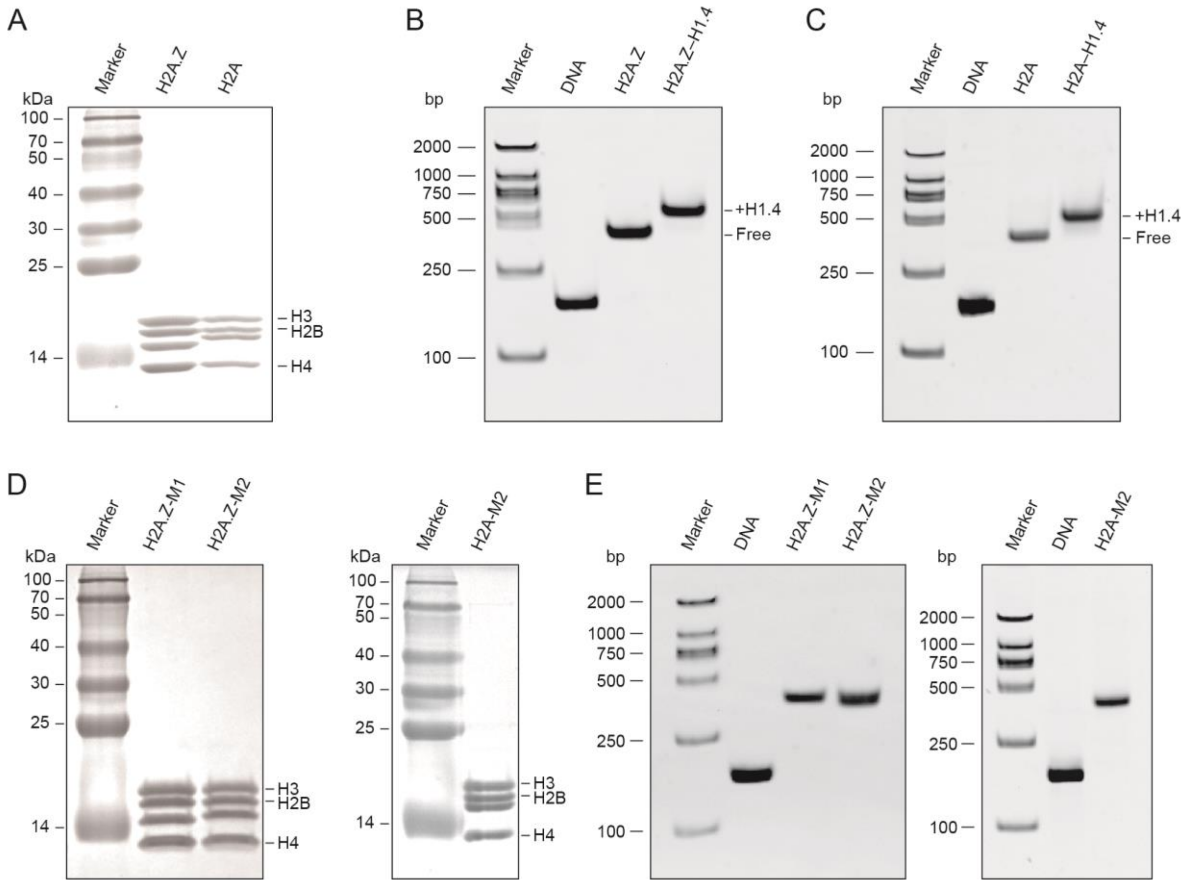


**Figure S1. Preparation and characterization of nucleosome samples.**

(A) SDS-PAGE analysis of reconstituted histone octamers containing H2A.Z or H2A. All core histone components are visible on an 18% SDS-PAGE.

(B) Native PAGE analysis of H2A.Z nucleosomes and H2A.Z–H1.4 complex. Samples were analyzed on a 5% native PAGE. Single bands indicate properly assembled particles.

(C) Native PAGE analysis of H2A nucleosomes and H2A–H1.4 complex. Samples were analyzed on a 5% native PAGE. Single bands indicate properly assembled particles.

(D) SDS-PAGE analysis of chimeric histone octamers H2A.Z-M1, H2A.Z-M2, and H2A-M2. Samples were analyzed on an 18% SDS-PAGE.

(E) Native PAGE analysis of chimeric nucleosomes H2A.Z-M1, H2A.Z-M2, and H2A-M2. Samples were analyzed on a 5% native PAGE. Single, uniform bands indicate homogeneous and stable assembly.

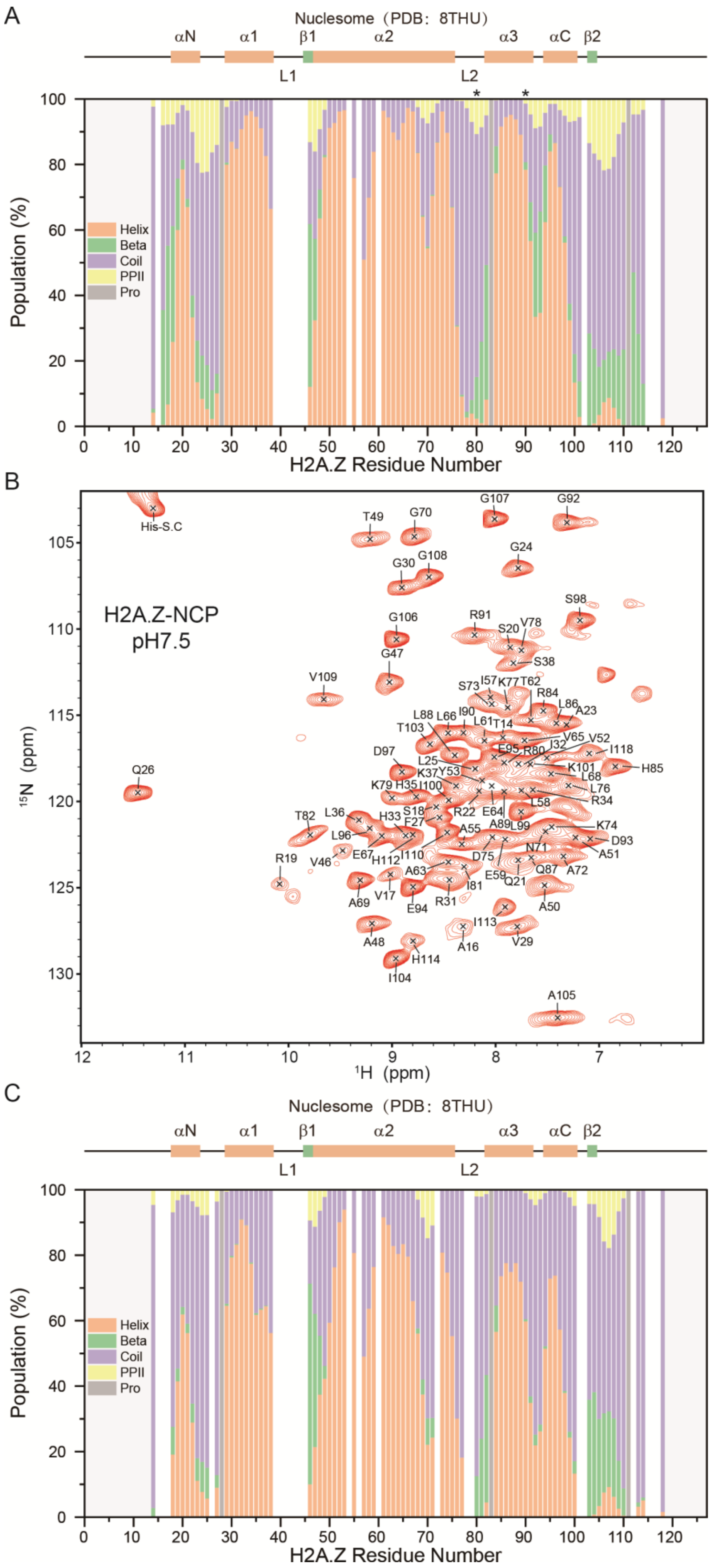


**Figure S2. Backbone resonance assignment and secondary structure analysis of H2A.Z.**

(A) Chemical shift-based secondary structure analysis (Δ2D) based on assigned H2A.Z chemical shifts at pH 6.5.

The cryo-EM-derived secondary structure (PDB: 8THU[10]) is shown at the top. Asterisks indicate residues with overlapping spectral peaks.

(B) Assigned 2D NH spectrum of H2A.Z at pH 7.5. A total of 87 out of 102 non-proline residues in H2A.Z (residues 14–118) were assigned, corresponding to 85% coverage of the rigid core.

(C) Chemical shift-based secondary structure analysis (Δ2D) based on assigned H2A.Z chemical shifts at pH 7.5. The cryo-EM-derived secondary structure (PDB: 8THU[10]) is shown at the top.

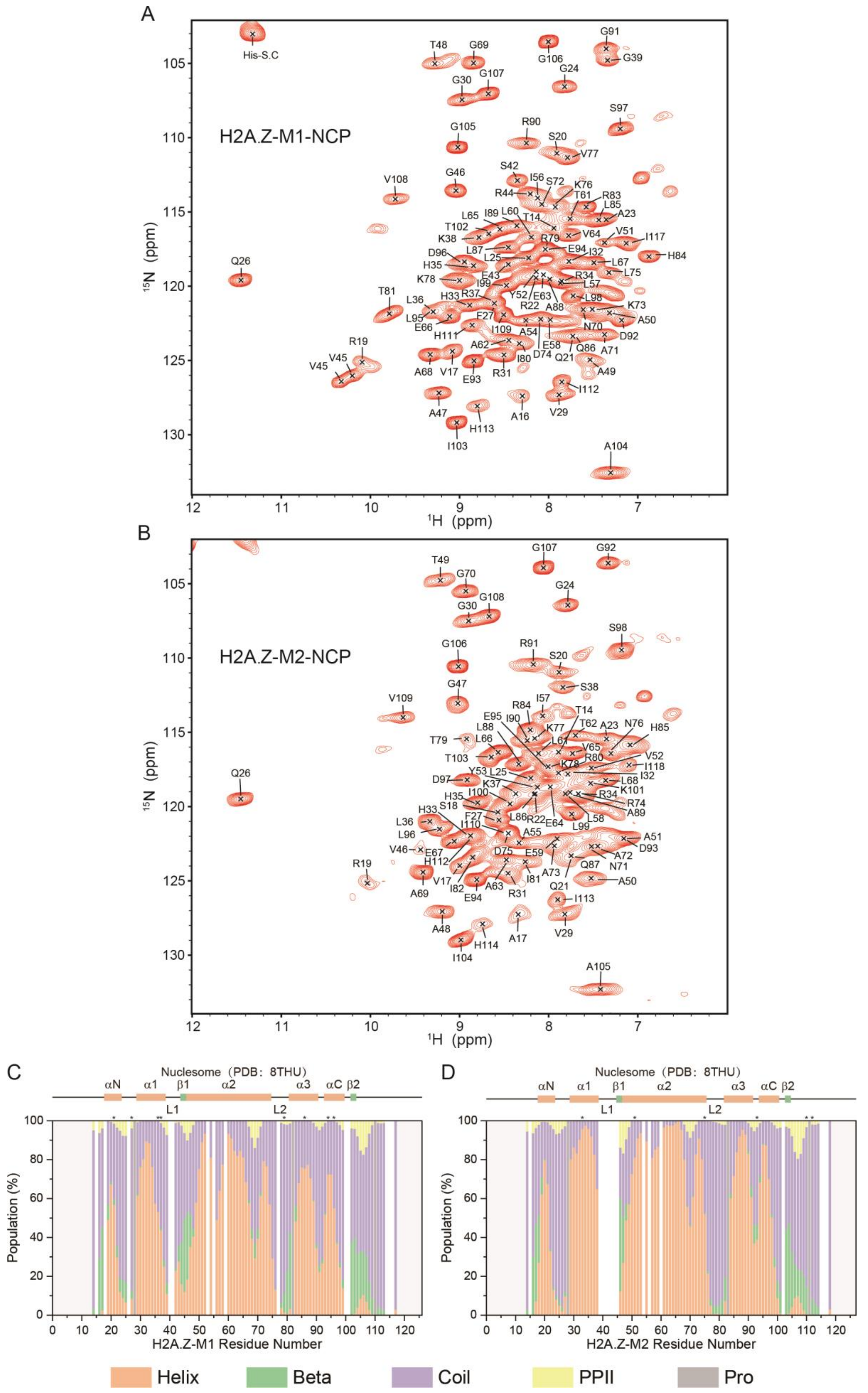


**Figure S3. Backbone resonance assignment and secondary structure analysis of H2A.Z chimeras.**

(A) Assigned 2D NH spectrum of H2A.Z-M1 at pH 7.5. A total of 89 out of 101 non-proline residues in H2A.Z-M1 (residues 14–117) were assigned, corresponding to 88% coverage of the rigid core.

(B) Assigned 2D NH spectrum of H2A.Z-M2 at pH 7.5. A total of 87 out of 102 non-proline residues in H2A.Z-M2 (residues 14–118) were assigned, corresponding to 85% coverage of the rigid core.

(C) Chemical shift-based secondary structure analysis (Δ2D) based on assigned H2A.Z-M1 chemical shifts at pH 7.5. The cryo-EM-derived secondary structure (PDB: 8THU[10]) is shown at the top.

(D) Chemical shift-based secondary structure analysis (Δ2D) based on assigned H2A.Z-M2 chemical shifts at pH 7.5. The cryo-EM-derived secondary structure (PDB: 8THU[10]) is shown at the top.

Asterisks in (C) and (D) indicate residues with overlapping spectral peaks.

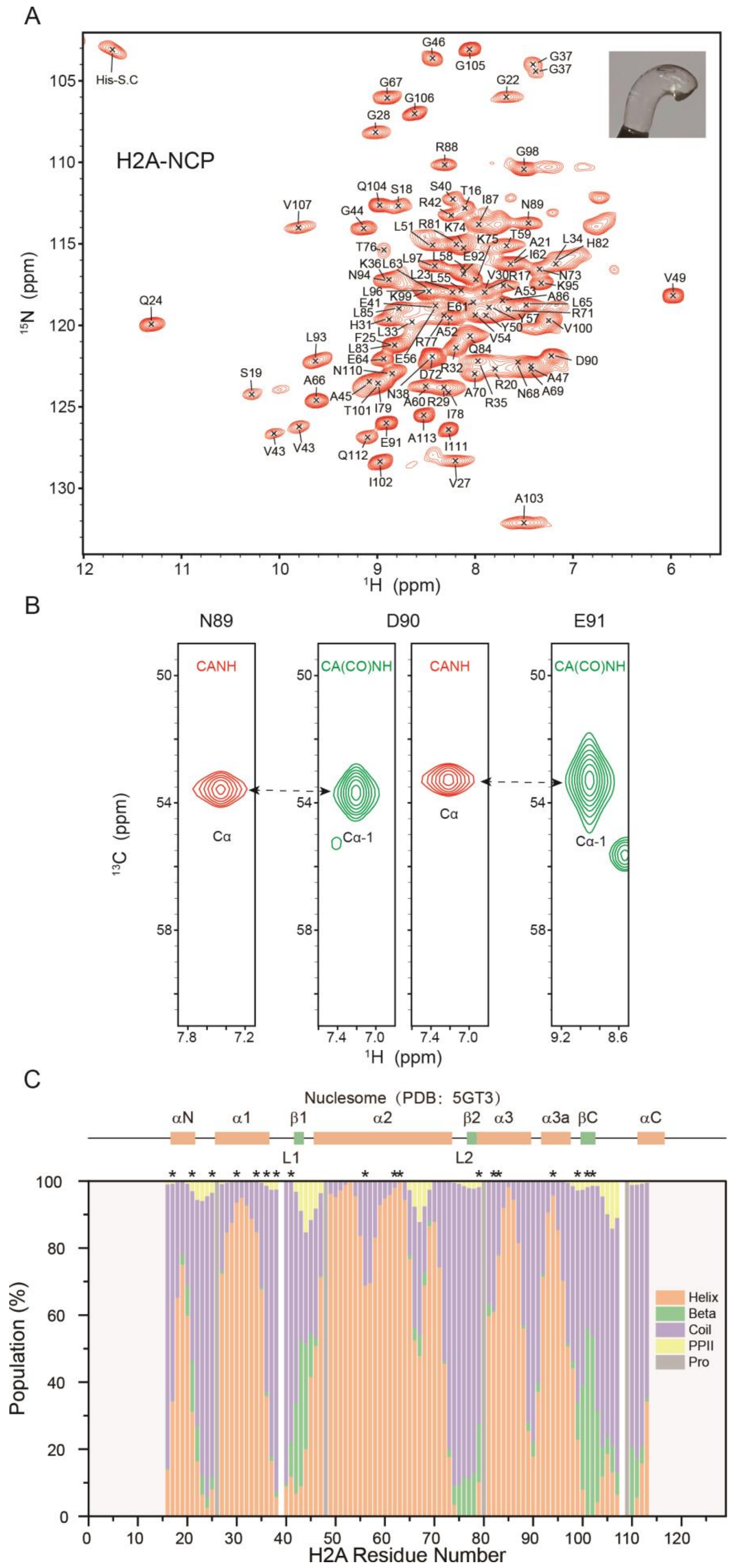


**Figure S4. Backbone resonance assignment and secondary structure analysis of H2A.**

(A) Assigned 2D NH spectrum of H2A at pH 7.5. A total of 92 out of 94 non-proline residues in H2A (residues 16–

113) were assigned, corresponding to 98% coverage of the rigid core. Inset: photograph of the gel-like sedimented nucleosome sample used for solid-state NMR experiments.

(B) Representative sequential walk from 3D CANH (red) and CA(CO)NH (green) spectra illustrating the sequential connectivity of residues N89→D90→E91 in H2A.

(C) Chemical shift-based secondary structure analysis (Δ2D) based on assigned H2A chemical shifts at pH 7.5. The X-ray-derived secondary structure (PDB: 5GT3[11]) is shown at the top. Asterisks indicate residues with overlapping spectral peaks.

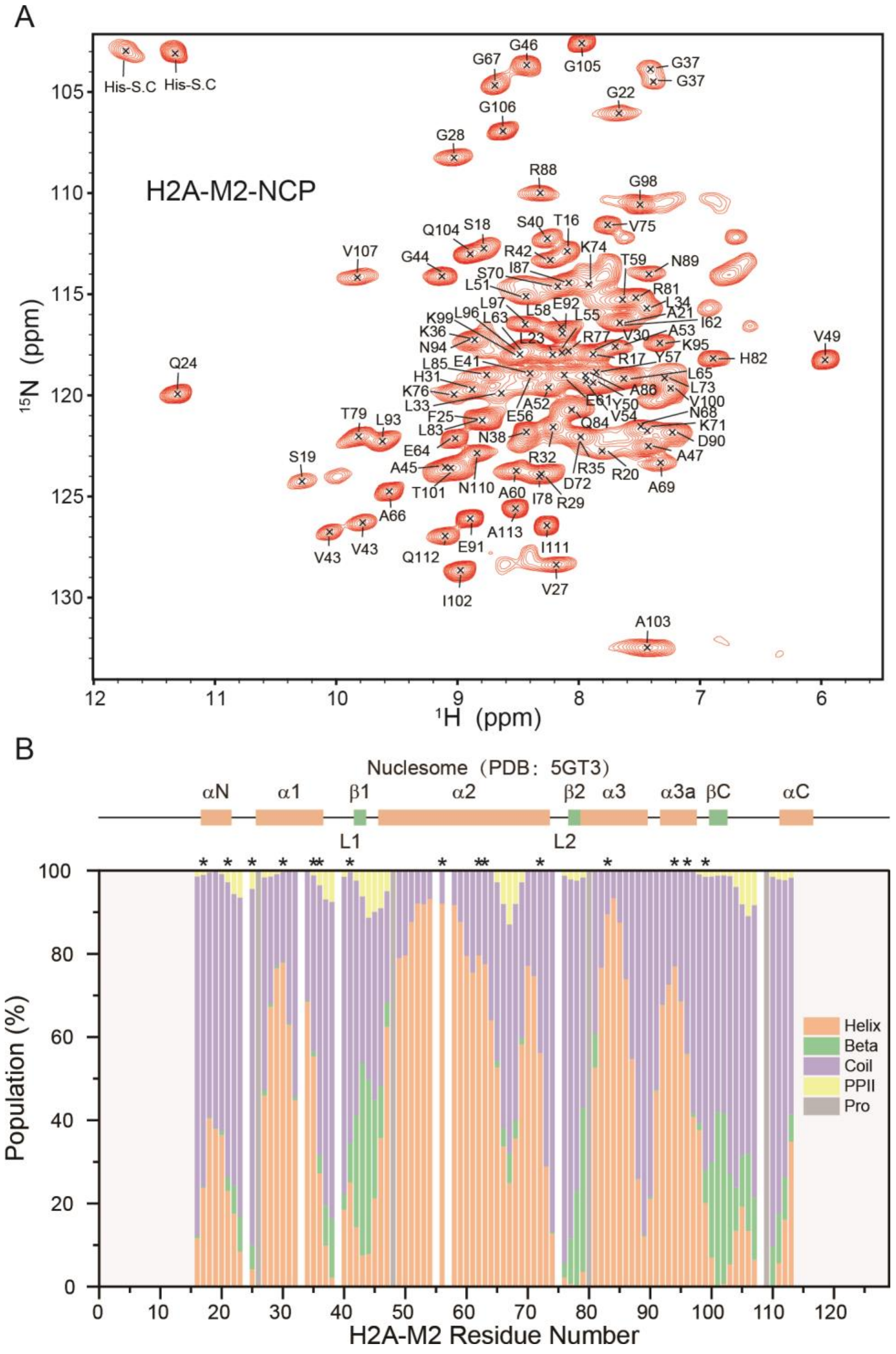


**Figure S5. Backbone resonance assignment and secondary structure analysis of the H2A-M2 chimera.**

(A) Assigned 2D NH spectrum of H2A-M2 at pH 7.5. A total of 92 out of 94 non-proline residues in H2A-M2 (residues 16–113) were assigned, corresponding to 98% coverage of the rigid core.

(B) Chemical shift-based secondary structure analysis (Δ2D) based on assigned H2A-M2 chemical shifts at pH 7.5. The X-ray-derived secondary structure (PDB: 5GT3[11]) is shown at the top. Asterisks indicate residues with overlapping spectral peaks.

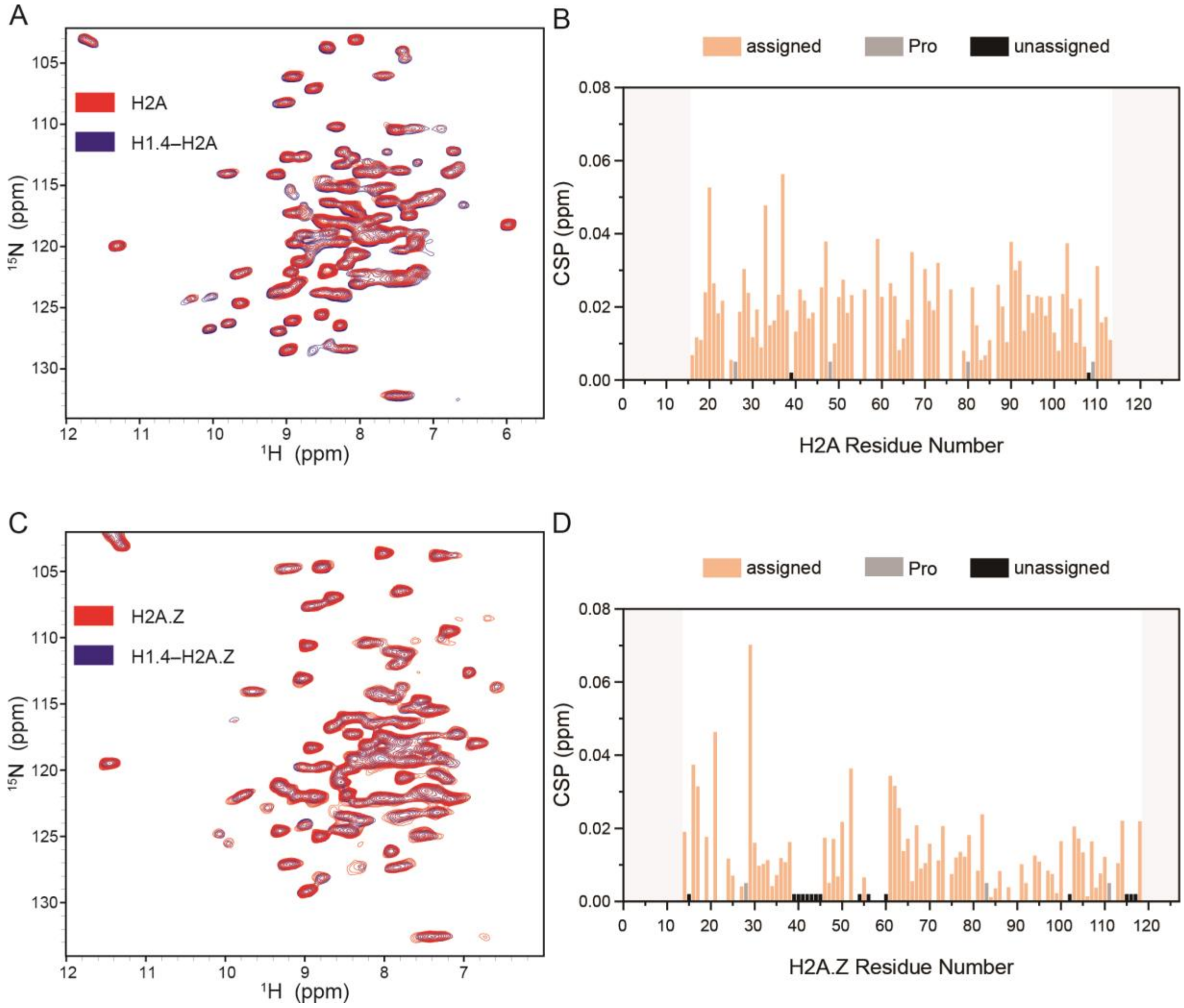


**Figure S6. H1.4 does not perturb the core structural domains of H2A or H2A.Z.**

(A) Overlay of cross-polarization-based 2D NH spectra of H2A in the absence (red) and presence (blue) of H1.4. The high spectral overlap indicates minimal conformational perturbation of the H2A core.

(B) Quantitative chemical shift perturbation (CSP) analysis comparing H2A and the H1.4–H2A complex. Most residues show negligible CSP values, consistent with H1.4 engaging linker DNA rather than the core histone fold.

(C) Overlay of cross-polarization-based 2D NH spectra of H2A.Z in the absence (red) and presence (blue) of H1.4. The high degree of spectral overlap indicates minimal conformational perturbation of the H2A.Z core.

(D) Quantitative CSP analysis comparing H2A.Z and the H1.4–H2A.Z complex. Most residues show negligible CSP values, consistent with H1.4 engaging linker DNA rather than the core histone fold.

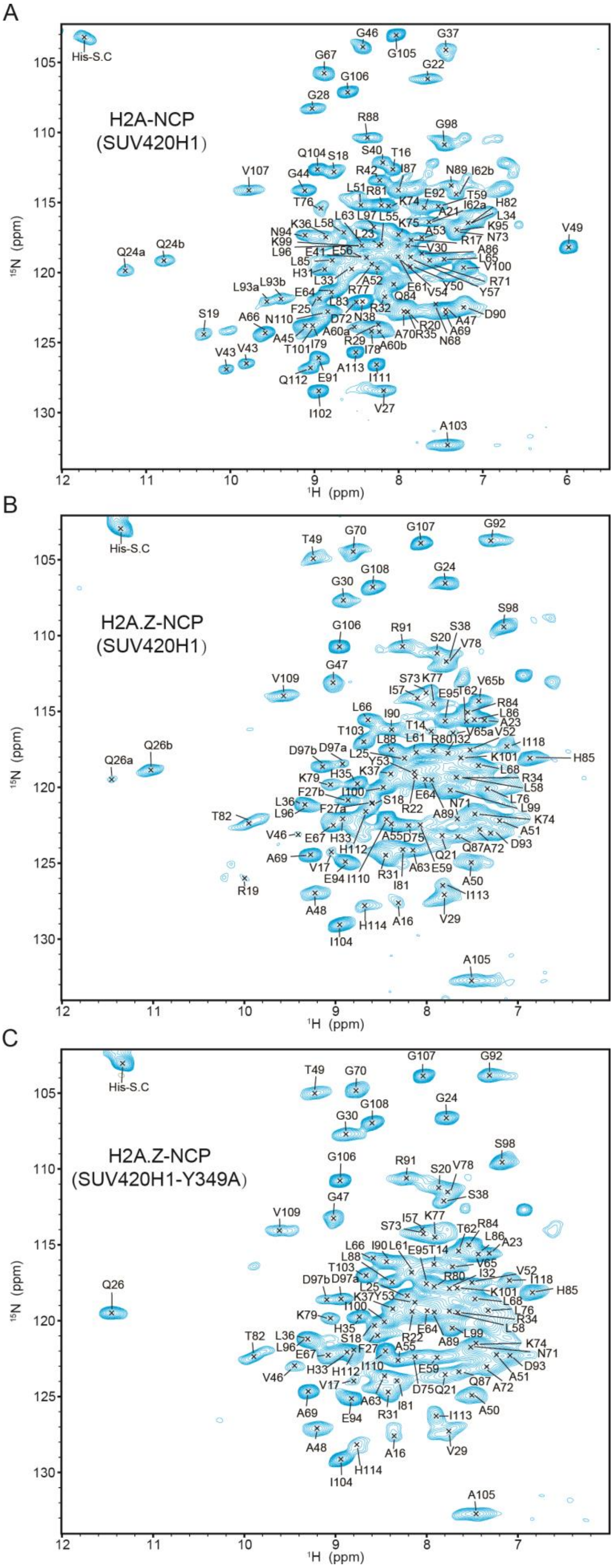


**Figure S7. Backbone resonance assignments of H2A and H2A.Z in SUV420H1–NCP[167] complexes.**

(A) Assigned 2D NH spectrum of H2A in the SUV420H1–H2A nucleosome complex at pH 7.5. A total of 92 out of 94 non-proline residues in H2A (residues 16–113) were assigned, corresponding to 98% coverage of the rigid core.

(B) Assigned 2D NH spectrum of H2A.Z in the SUV420H1–H2A.Z nucleosome complex at pH 7.5. A total of 87 out of 102 non-proline residues in H2A.Z (residues 14–118) were assigned, corresponding to 85% coverage of the rigid core.

(C) Assigned 2D NH spectrum of H2A.Z in the SUV420H1-Y349A–H2A.Z nucleosome complex at pH 7.5. A total of 87 out of 102 non-proline residues in H2A.Z (residues 14–118) were assigned, corresponding to 85% coverage of the rigid core.

For some residues, the peak corresponding to the free nucleosome state is labeled "a", and the additional peak that appears in the complex, representing a second population arising from SUV420H1 binding, is labeled "b".

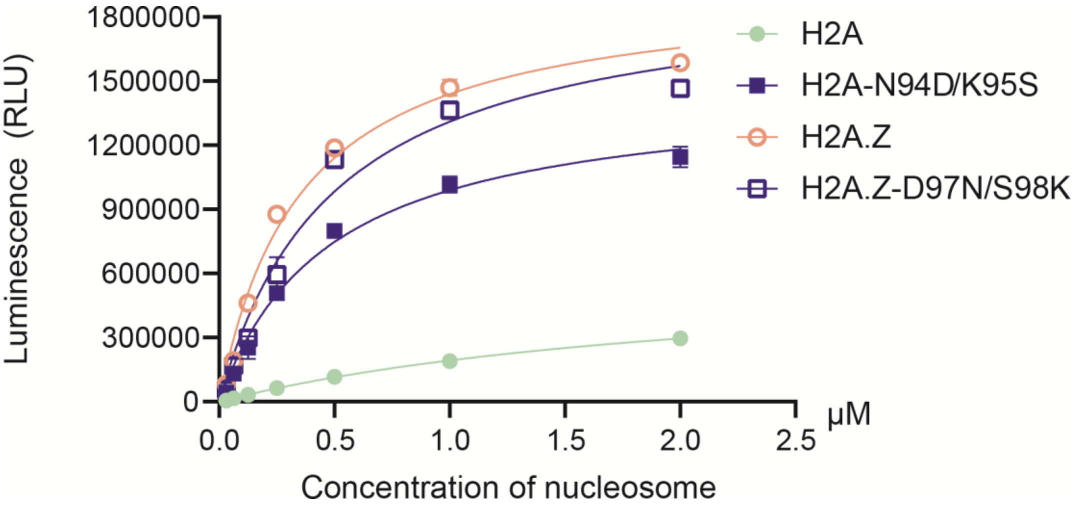


**Figure S8. Bidirectional exchange of the DS and NK motifs did not fully reverse SUV420H1 activity levels between H2A.Z and H2A.** MTase-Glo analysis results of HMT assays carried out with wild-type SUV420H1 on nucleosomes containing H2A, H2A-N94D/K95S, H2A.Z, or H2A.Z-D97N/S98K are shown. The results show that reciprocal mutation of the DS and NK sites does not fully interchange the SUV420H1 activity of H2A.Z and H2A, indicating that determinants beyond the DS motif contribute to SUV420H1's preferential recognition of H2A.Z.

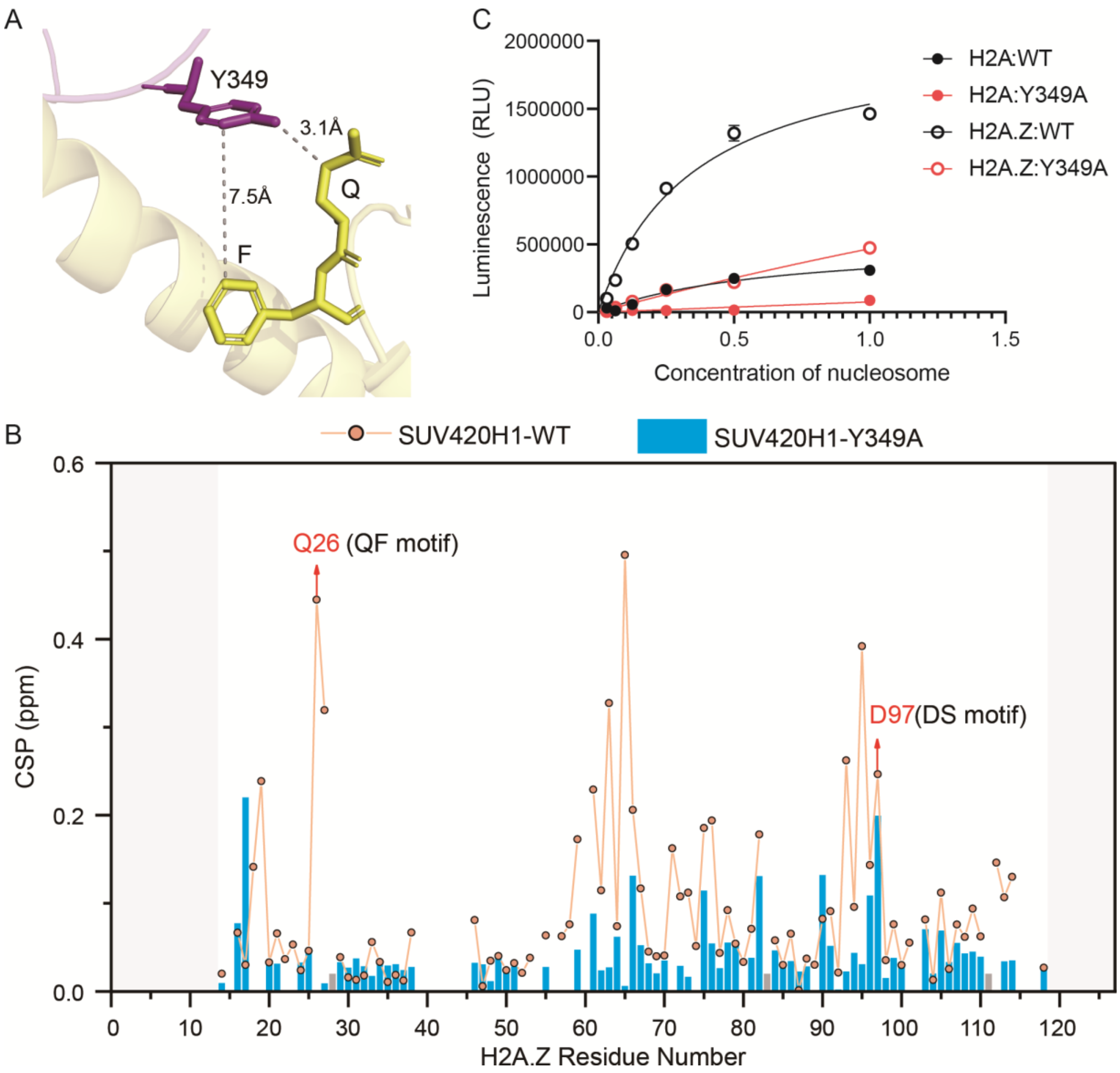


**Figure S9. Residue Y349 in SUV420H1 mediates nucleosome binding and catalytic activity.**

(A) Structural context of Y349 and the conserved QF motif in the SUV420H1–nucleosome complex (PDB:7YRG[12]). The dashed line indicates the distance between the Y349 side chain and the QF motif.

(B) Chemical shift perturbation (CSP) analysis of wild-type and the Y349A mutant SUV420H1 upon binding to H2A.Z nucleosomes. The Y349A mutation attenuates CSP across most residues, indicating impaired enzyme–nucleosome interaction.

(C) The MTase-Glo analysis results of the HMT assay carried out using wild-type SUV420H1 (WT) or Y349A mutant on nucleosomes containing H2A or H2A.Z. The Y349A mutation substantially reduces SUV420H1 catalytic activity, establishing Y349 as critical for enzymatic activity.
